# Image-based ecological assessment of deep-sea sponge, coral and other cnidarian assemblages through a morpho-functional approach

**DOI:** 10.1101/2025.08.18.670867

**Authors:** Mélissa Hanafi-Portier, Sarah Samadi, Paco Cárdenas, Eric Pante, Karine Olu

## Abstract

Porifera and Cnidaria are key phyla that contribute to the structural complexity and functioning of marine ecosystems and are commonly considered in the identification of Vulnerable Marine Ecosystems. Their diversity and distribution largely rely on seafloor images, but low-resolution taxonomic identification is hindered by morphological plasticity and convergence, and by the need for microscopic diagnostic characters or molecular markers that are not accessible from images. Higher-rank taxonomy also poorly reflects their morphological and functional diversity. To address this issue, we developed an image-based classification combining multiple macro-morphological traits to define statistically derived morpho-functional groups (MFGs) which were then used as response variables in ecological analyses instead of taxa. This framework provides a reproducible and transferable approach for comparing patterns of habitat-forming taxa across regions, regardless of the level of taxonomic knowledge available, by relying on observable morphological traits rather than uncertain taxonomic IDs. We applied this framework to sponge and cnidarian assemblages from seamounts and island slopes in the Mozambique Channel, a poorly sampled region where taxonomic limitations are acute. We assessed the relevance of an MFG approach to provide additional information on sponge and cnidarian assemblage diversity, spatial variability and relationships to environmental gradient, as well as their structuring roles on the surrounding megabenthic community, compared with the available taxonomic resolution. We identified 37 sponge and 34 cnidarian MFGs revealing substantial morphological variation within the coarse taxonomic categories available from imagery. MFGs captured greater spatial differentiation and local variability than the taxonomic classification, particularly for sponges, for which only two taxonomic classes were available. Several MFGs were also associated with particular environmental conditions and with other megabenthic taxa, although the proportion of community variation explained by these relationships was low. Overall, the MFG approach, especially effective for sponges, allowed to retain and quantify morphological and structural information of habitat-forming taxa that can be lost using coarse image-based taxonomy. Its multi-trait classification can be readily translated into existing hierarchical image-classification systems such as CATAMI, allowing finer morphological information to be retained while maintaining consistency with broader standardised categories. Finally, the framework proposes metrics to detect and quantify the heterogeneity and spatial variability of potentially vulnerable biogenic habitats in imagery-based studies.

## Introduction

In marine benthic environments, sessile, filter-feeding organisms may form large aggregations to which a community of associated species can be found. To unify the vague and often overlapping terminology used to describe these habitats (such as “cold-water coral reefs”, “coral/sponge gardens”, or “sponge grounds”), Rossi & Bramanti (2020) recently introduced the overarching concept of Marine Animal Forests. This concept groups various epi-zoobenthic taxa that share a common trait, i.e., a three-dimensional architecture extending from a few centimetres to a couple of meters in height. Much like trees in terrestrial forests, these biogenic structures provide essential ecosystem functions. Among such benthic organisms, sponges and cnidarians can form large structures (metre scale), which sustain many functional roles like providing shelter and nurseries for other species or regulating biogeochemical/nutrient cycles (Bell, 2008; Rossi et al., 2017). Through their three-dimensional structure, these habitat-forming taxa contribute to habitat complexity, which supports high levels of diversity of the associated community (Bell, 2008; Quattrini et al., 2012; Beazley et al., 2013; Rogers et al., 2014; Hawkes et al., 2019). Their 3D-structure also interacts with currents and locally modulates the current regime, creating turbulence and resuspension retaining food particles or preys for their associated fauna (Buhl-Mortensen et al., 2010).

In deep-sea environments, seamounts are places where habitat-forming taxa such as coral and sponge may create marine animal forest. Seamounts are targeted in some regions by industrial fishing and are also potential targets for deep-sea mining of rare metals (e.g., cobalt-rich crust) that may cover their surfaces (Hein et al., 2010). Such activities can have adverse consequences – razing activities, smothering – on the benthic suspension feeder communities that commonly occur over these topographic features (Baco et al., 2020; Morgan & Baco, 2021). In addition to their structuring role as habitat forming organisms, it is assumed that sponges and cnidarians living in the deep sea are slow-growing and long-lived taxa, and are structurally fragile, making them poorly resilient to mechanical disturbances from anthropic activities. Therefore, they are defined as indicator taxa of Vulnerable Marine Ecosystems (VME; FAO, 2009; Williams et al., 2020a; Watling & Auster, 2021). However, little is known about the factors that structure the distribution of these VME’s indicator taxa and how they structure the diversity of the associated fauna.

The analysis of their distribution, especially at small-scales, and of their role in the structure of deep-sea communities requires deployment of submarine imaging systems. However, a major source of difficulty remains the taxonomic ID of these taxa from images. Identification of sponges and cnidarians from bottom images is hardly achievable beyond the class or order levels because in the poorly explored deep-sea environment, the species inventory is very incomplete and no images-based identifications keys can be established. Consequently, the identification of species only relies on taxonomic diagnostic characters that are not visible on images (e.g., spicules and fibers for sponges, sclerites for cnidarians) (Hooper & van Soest, 2002; Pante et al., 2012b). Given this limitation, many ecological studies based on images use any observable features (such as colour or global shape) to define morphotypes as proxy of species entities outside the formal framework of taxonomy. Such approaches make it difficult to compare data among regions and studies. For these taxa, morphology is highly variable both across higher taxonomic levels and within species, and both at macro and microscopic scales (Bell, 2007; Todd, 2008). This variability is due to intraspecific polymorphism, phenotypic plasticity, and morphological convergences among distant species. It is also worth noting that the current state of the alpha taxonomy for these groups remains very incomplete, and frequently challenged by molecular data (e.g., McFadden et al., 2022). Consequently, the use of morphotypes to approximate taxonomic diversity remains questionable (Krell, 2004). While some studies revealed a positive correlation between morphological and taxonomic diversity (Bell & Barnes, 2001; Bell & Barnes, 2002; Hadi et al., 2015), there is no consensus regarding these relations (e.g., Pica et al., 2018; Shaffer et al., 2019) nor with the functional diversity (Bell, 2007).

Therefore, taxonomic ID of these groups from images remains a weak approach to study the diversity and functional role of sponges and cnidarians in benthic communities. To overcome this difficulty, some authors suggested to analyse directly macro-morphologies to determine how it varies in response to the environmental factors (Denis et al., 2017; Mary George et al., 2018; Zawada et al., 2019a, 2019b, Schönberg, 2021) and whether their variability structure the associated fauna (e.g., Buhl-Mortensen et al., 2010, 2016). Studies, mostly on coral reef ecosystems and shallow waters, have revealed variability of the external morphology of these sessile fauna in response to a range of environmental parameters, including light (Wilkinson & Vacelet, 1979; Chappell, 1980), sedimentation (Chappell, 1980; Bell et al., 2016; Pineda et al., 2016; Schönberg, 2016), hydrodynamics (Chappell, 1980; Pronzato et al., 1998; Bell & Barnes, 2000; Barnes & Bell, 2002; McDonald et al., 2002; Meroz-Fine et al., 2005; Paz-García et al., 2015; Schönberg, 2021), depth (Bell & Barnes, 2000; Barnes & Bell, 2002; Soto et al., 2018; Gökalp et al., 2020; Kramer et al., 2020), or substrate (Barnes & Bell, 2002; Duckworth, 2016). Such approaches, based on multiple traits quantifying the morphological diversity and structural complexity of biogenic habitats, were for example recently used to understand how coral reefs respond to anthropogenic activity (Zawada et al., 2019b) and to a bleaching event (Denis et al., 2017). To assess the role of sponge and cnidarian 3D-architectures on benthic communities, the review of Buhl-Mortensen et al. (2010) suggested the use of descriptors such as size, volume, complexity (surface/volume), branching patterns or flexibility of biotic structures, which have been reported to have an influence on the diversity of physically associated species on these 3D-structures, and to directly or indirectly benefit the megabenthic communities. The use of morphological descriptors (traits) therefore appears to be a potential functional predictor of the response (to environmental drivers) or effect (on communities/ecosystems) of sponges and cnidarians.

Owing to the difficulty in identifying these taxa from visual surveys, recent initiatives aiming to standardise the classification of habitat-forming taxa morphologies have been developed (e.g., SmarTarID, Howell et al., 2019). Schönberg (2021) has recently developed a standard classification to characterise sponges based on their functional morphologies rather than their taxonomic identities, adapted in particular to the use of images. This classification has adapted the CATAMI standards (Althaus et al., 2015), which proposes a mix between morphological and taxonomic classification to identify morphotypes for different taxonomic groups.

From this background literature and based on existing classifications (Althaus et al., 2015; Schönberg, 2021), we developed a classification of sponges and cnidarians adapted to a dataset derived from *in-situ* deep-sea imagery. Rather than assigning organisms to predefined morphological categories, such as those used in CAABs, our approach relies on multiple observable morphological traits and is mostly independent of taxonomic knowledge. Each unique combination of traits defines a preliminary morphotype. Morphotypes are then grouped into broader morpho-functional groups (MFGs) (*sensu* “morphology-based functional groups” in Kruk et al., 2010, 2011; see also Tsakalos et al., 2019) through numerical hierarchical classification based on their morpho-traits similarity. For both sponges and cnidarians, taxonomic ID is not used as a response variable. Instead, community analyses are conducted using MFGs rather than taxonomic or OTU/morphotype-based entities (species proxy).

Using MFGs, we hypothesised that part of the morphological structure described represents a functional structure (i.e., morphotypes grouped within an MFG share similar functional traits) and the traits selected have functional relevance in the ecosystem. We tested this methodology on a dataset from seamounts and island slopes in the Mozambique Channel which have already been analysed at the community level including a low taxonomic resolution for sponges and cnidarians and revealing patterns of spatial variability (Hanafi-Portier et al., 2024). However, these IDs reflect neither their taxonomic diversity nor the 3D-complexity observable on images and consequently underestimated their richness and the diversity of their morphologies.

Here, we compared taxonomic-based patterns for sponges and cnidarians with morpho-functional (morphological-based) ones to assess the benefit of the MFG approach in the assessment of (1) sponges and cnidarians diversity and spatial patterns at multiscales, (2) the environmental drivers shaping their morphological spatial structure, and (3) their potential structuring role on associated megabenthic communities.

## Material and Methods

### Study area and field acquisition

The data were obtained in the scope of the PAMELA project (Bourillet et al., 2013) during the campaigns PAMELA MOZ01 (Olu, 2014) on board the research vessel (R/V) *L’Atalante*, and PAMELA MOZ04 (Jouet & Deville, 2015) on board the R/V *Pourquoi Pas ?*. These cruises allowed the exploration of four seamounts (Glorieuses and Sakalaves platform, Jaguar and Hall Banks) and the terrace of Bassas da India island slope (**Figure 1**). The data were also obtained on three outer slopes (East, North, and West) of Mayotte Island, during the BIOMAGLO campaign (Corbari et al., 2017) on board the R/V *L’Antea* (**Figure 1**).

**Figure 1.**
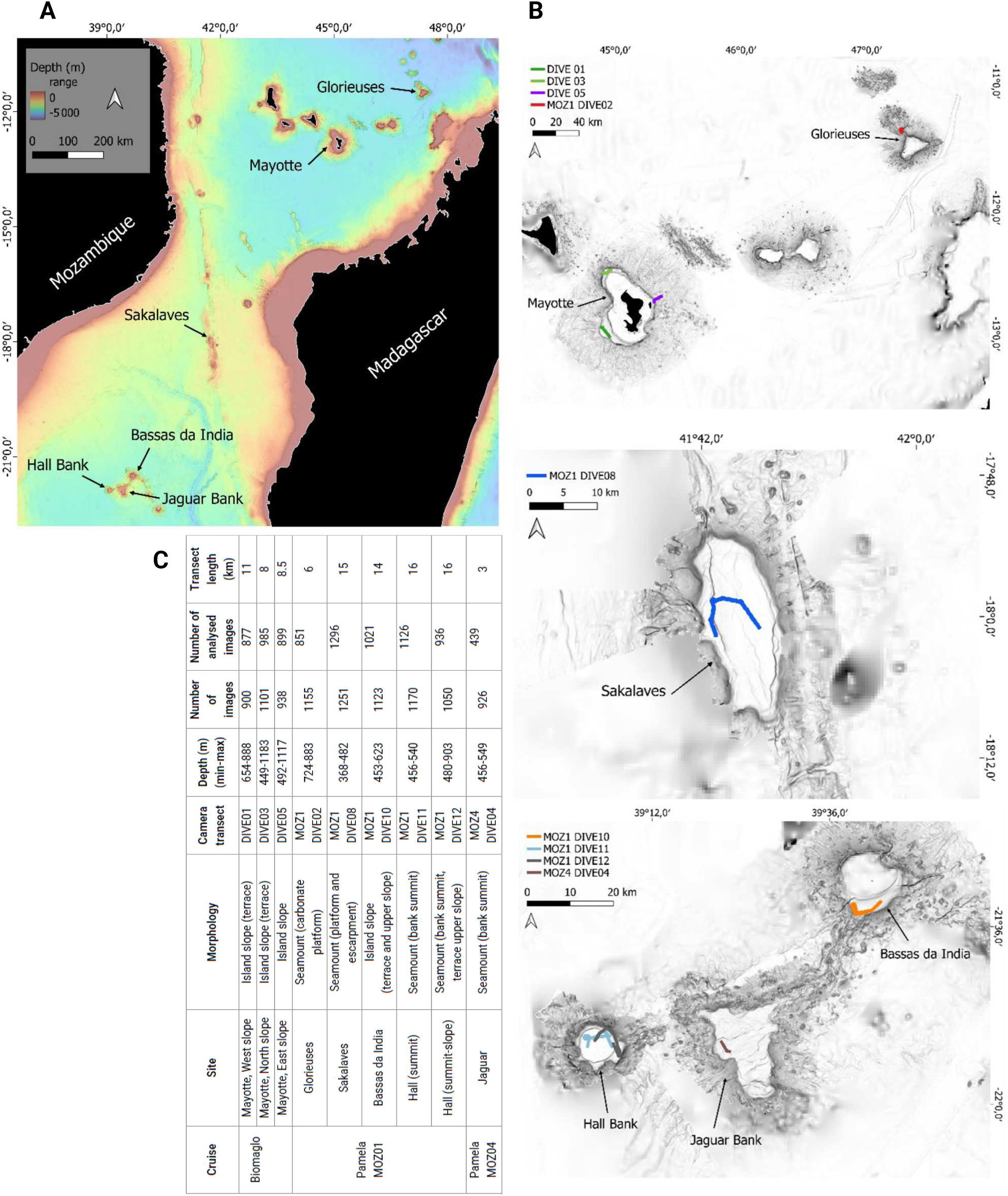
(A) Location of the different seamounts (Glorieuses and Sakalaves platforms, Jaguar and Hall Banks) and volcanic island external slopes (Mayotte, North, East and West slopes; Bassas da India southern slope) explored along the Mozambique Channel. (B) Location of the towed-camera transects within each site. (C) Details on sampling effort, location and depth for the nine towed-camera transects. Bathymetric grid from GEBCO Compilation Group (2023).

Images were captured from towed-camera transects (SCAMPI, French IFREMER fleet) in a vertical position at intervals of 30 s (except for Glorieuses, ∼15 s), at an altitude of 2.5-3 m and a speed of 0.5 m/s with a NIKON D700 HD camera (focal length 18 mm, resolution 4 256 × 2 832 pixels). The images were georeferenced by a positioning system attached to the camera, using the ADELIE application developed at IFREMER and implemented using ArcGiS V10.3 software. High resolution bathymetry and acoustic reflectivity data were acquired with a multibeam echosounder (Kongsberg EM122 for the deepest areas and EM710 for the shallowest areas) (Audru et al., 2006; Courgeon et al., 2016). CTD (temperature, salinity) and oxygen data were acquired during the Scampi dives from CTD and optode (MicroCAT, Seabird) sensors mounted on the camera frame. More specifics on the study area and field acquisition are available in the Material and Methods section in Hanafi-Portier et al. (2024).

A total of nine camera transects were processed for this study (**Table 1**). All nine transects were used for sponge classification while five transects from the northern area (Glorieuses seamount and the three outer slopes of Mayotte) were used for cnidarians due to data traceability concerns. Indeed, the cnidarian morphotypes for sites south of the Mozambique Channel were annotated manually (without the use of annotation software at the time) by another observer. To avoid biasing the data set due to inconsistencies in observations and difficulties in locating morphotypes within the images for data verification and standardising annotations, we have retained only the data from the seamounts north of the Channel for cnidarians. Prior to annotation, an image quality filter was applied such as all images with high motion blur, too dark, or an altitude below 2 m or exceeding 6 m above the substratum were discarded to ensure optimal visibility of the features.

**Table 1.** Morphological traits used for the classification of sponges in morpho-functional groups.

| Traits | Modalities | Description |
| --- | --- | --- |
| Sieve plates | 0/1 | Presence of sieves plates perforated by atrial cavities that cover the osculum. E.g., Euplectellidae. |
| Large spicules (Hispidity) | 0/1 | Large visible spicules that cover the body of the sponge giving a hairy appearance. E.g., Pheronema. |
| Internal cavities | 0/1 | Presence of cavities (visible holes, spaces between sheet-like or holes within branche-like features) as potential refuge for associated fauna. |
| Branches/outgrowths | 0/1 | Structures with outgrowths, reminiscent of branches. |
| Compactness | Aerial | Delicate, thin, appears "fragile"-like structure, having few seabed surface contact, but high seawater surface contact. |
|  | Intermediate | Some openings in the structure. Intermediate in their volume homogeneity and surface contact. |
|  | Compact | A compact and homogeneous volume, with little or no airspace/opening, with high seabed surface contact. |
| Osculum structure and visibility | Single and large and visible | A large single exhalant opening (osculum) on the top of the sponge, $\geq 50\%$ of the sponge's body. |
|  | Small and visible | Single or multiple medium to small oscula on the sponge's body. |
|  | Not visible | Either too small to be observed on images, or closed according to water stream conditions, or unfavourable viewing angle. |
| Body size | Small | 2-5 cm |
|  | Medium to large | >5-15 cm |
|  | Very large | >>15 cm |
| Thinly encrusted (Crust) (CAAB 10 000923) | 0/1 | Forming a thin crust (flat sheet) over a hard substrate (very low profile) (Schönberg, 2021). |
| Thickly encrusted (Crust) (CAAB 10 000924) | 0/1 | Forming a thick crust (few mm to ~1 cm), slightly elevated from the hard substrate (low profile) (Schönberg, 2021). |
| Creeping (Crust) (CAAB 10 000917) | 0/1 | Growth following the substrate (repent sponge), slightly thick in width, and elevated from the substrate. Some morphologies can be curved (incomplete vase shape, not closed). |
| Ball (Massive) (CAAB 10 000905) | 0/1 | Globular/ball shaped sponges: can be small or large, compact or with a fragile appearance. |
| Amorphous simple (Massive) (CAAB 10 000904) | 0/1 | Irregularly shaped sponges, with no particular morphology or detectable structures (e.g., branches), generally quite large. |
| Amorphous composite (Massive) (CAAB 10 000925) | 0/1 | Irregularly-shaped sponges, composed of merged sub-units (Schönberg, 2021). |
| Fistular/Cryptic (Erect) (CAAB 10 000908) | 0/1 | The sponge is partly buried in the sediment, with only syphons emerging from the substrate. |
| Columnar (Erect) (CAAB 10 00916) | 0/1 | Simple erect sponge, higher than wide, not branched, solid in cross-section (Schönberg, 2021). |
| Laminar (Erect) (CAAB 10 000913) | 0/1 | Erect growth forms with flattened body, can have a small stalk (have a low anchoring area) (include fans, blades and spatulas) (Schönberg, 2021). |
| Stalked (Erect) (CAAB 10 000906) | 0/1 | Include all erect forms, standing up from the substrate from a stalk, small or high. Upper part can be cup, ball, branching-shaped, etc., but the stem is inactive, meaning it does not have oscules/ostiae (Schönberg, 2021). |
| Bushy/arborescent (Erect) (CAAB 10 000915) | 0/1 | Dense 3D-structure, forming a little bush/tree and narrow interstice between extended structures (e.g., "branches"). Oscules/ostia scattered across the sponges (Schönberg, 2021). |
| Amphoras (Narrow cup-like) (CAAB 10 000927) | 0/1 | Hollow with a narrow opening (Schönberg, 2021). |
| Tube (Narrow cup-like) (CAAB 10 000911) | 0/1 | Hollow along most of their body. Apical opening of similar diameter as the body (Schönberg, 2021). |
| Vase-shaped (~Wide cup-like) | 0/1 | Deep and wide-opened osculum, body hollowed in cross-section with apical part having the largest diameter. |
| Cup (Wide cup-like) (CAAB 10 000919) | 0/1 | Complete wide cup, concave with apical part having the largest diameter (Schönberg, 2021). |
| Incomplete cup (Cup-like) (CAAB 10 000918) | 0/1 | Intermediate forms among erect-laminar sponges, cups and tables. Semi-closed (Schönberg, 2021). |

### Megabenthic community and morpho-functional groups processing

#### Megabenthic community dataset

The megabenthic community dataset comprised 99 taxa of sessile and mobile invertebrates, and fish, of various taxonomic resolutions, identified from 9 614 images, using the online image annotation platform BIIGLE 2.0 (Langenkämper et al., 2017). Details of the method for identifying and classifying megafauna in images are described in Hanafi-Portier et al. (2021). Details of the method for selection and curation of the megabenthic community dataset reused in the present study, are described in Hanafi-Portier et al. (2024). Faunal densities were standardised in 200 m^2^ (∼60 m linear) sample units, to capture sufficient faunal density while conserving the scale of habitat heterogeneity. This megabenthic dataset based on taxonomic identifications was reused for comparison with the one using MFGs of: (1) cnidarians and sponges community metrics (richness, beta diversity) by analysing each one separately, (2) the role of environmental variables on their spatial structure and (3) their role in structuring the rest of the megabenthic community, compared to the environmental variables. In the paper, we will refer to the term ‘associate’ to describe megabenthic taxa either directly physically associated with sponge and cnidarian MFGs, or closely co-occurring in the same area. Based on the definition of the word ‘coral’ by Cairn (2007), we will refer to the term ‘cnidarians’ in order to include corals and other cnidarians (mainly Actiniaria).

#### Sponge and cnidarian primary annotations into morphotypes

For the image analysis of sponges and cnidarians, we built upon the primary annotations established by Hanafi-Portier et al. (2024). The existing annotations were systematically re-classified using the ’Largo’ tool within the BIIGLE interface.

We classified the images of sponges and cnidarians into homogeneous morphotypes according to criteria observable in the images: general shape (e.g., ball, tube, amorphous), other structures (e.g., long spicules giving a hairy appearance), and colour. We then assigned each morphotype to a size category (small, medium, large) using the BIIGLE measuring tool (see size classes in **Supplem. mat. 1 and Supplem. mat. 2**). In some cases, a morphotype corresponded to a single taxa identified at low taxonomic ranks (e.g., genera - *Aphrocallistes* for sponges; *Umbellula*, *Bathyalcyon*, *Enallopsammia* for cnidarians; or families - Euplectellidae, Hyalonematidae for sponges; Primnoidae for cnidarians), because it corresponds to a unique morphology easily recognizable from images. However, most of the taxonomic IDs, achieved at the class (within sponges) and order (within cnidarians) ranks, were divided into several morphotypes. The use of the BIIGLE ‘Largo’ option, which allows to visualise and review the already classified morphotypes through an image catalogue, facilitated the classification of the remaining images. Unclassifiable morphologies (e.g., from blurry images) were discarded from the analyses. A total of 24 835 annotations of sponges and 4 005 annotations of cnidarians were classified into morphotypes.

#### Morphological traits selection and definition

The morphotypes were then formally described through a set of morphological traits. Both for sponges and cnidarians, the morphological traits were based on the CATAMI catalogue, by Althaus et al. (2015), whose classification of sponge morphologies has recently been reviewed with the framework of Schönberg, 2021. In this scheme, each morphology has a unique CAAB – Codes for Australian Aquatic Biota – number.

In Schönberg (2021) framework, the sponge morphologies are classified based on their inferred ecological function, using a hierarchical structure, and allows scoring at different levels of resolution (ranging from 4 to 21 functional morphologies) according to the scientific purpose. To assess environmental conditions of sponges, at least 14 morphologies are recommended to be scored. We followed this classification, and scored 16 morphologies, as we differentiated a category ‘vase’ as an intermediate form among complete wide cups (CAAB 10 000919), narrow cups (tubes and chimneys) (‘CAAB 10 000911’) and barrels (CAAB 10 000907), considering vasiform sponge morphotypes observed on the images would respond differently to sedimentation than the other wide-cups morphologies observed (**Table 1**).

For cnidarians, the CATAMI classification relies on a combination of high taxonomic ID ranks as a first entry in their classification tree (e.g., orders Scleractiniaria, and Antipatharia; class Octocorallia) with morphological categories, and is particularly relevant to characterise elements such as robustness (fleshy/hard), complexity (single/multiple axis). In our approach, we did not consider the taxonomy of corals as a first entry in the classification. Our rationale was to dissociate, as much as possible, morpho-functional diversity from taxonomic diversity, however, some traits still required basic knowledge on coral taxonomy (e.g., anchoring mode, flexibility) at high rank. We have used and declined the proposed categories into multiple traits. For example, the ‘CAAB 11 168906’ corresponds to the traits combination ‘Black & Octocorals > Branching (3D) > Non-fleshy > Bottle-brush > Simple (axe)’. We declined this unique pathway into a set of different combination of traits: e.g., growth form (e.g., bottle-brush), presence of branches, dimension (2D/3D), skeleton flexibility that we declined into four sub-categories (fleshy/semi-fleshy/semi-rigid/rigid), and number of axes that we declined into three sub-categories (single/multiple/no main axis) (**Table 2**).

**Table 2.** Morphological traits used for the classification of cnidarians in morpho-functional groups.

| Traits | Modalities | Description |
| --- | --- | --- |
| Dimension | 2D (planar) | Structure in a two-dimensional plane ("flattenable" structure). E.g., Antipatharia, Plexauridae) |
|  | 3D | Three-dimensional (a volume). E.g., Chrysogorgia |
| Modularity | Solitary | Single polyp within the same skeleton (one individual). E.g., Javania sp. |
|  | Colonial | Several polyps within the same skeleton. E.g., Enallopsammia |
| Anchoring | Calcified disc | A calcified disc attached to rock bottom (holdfast). E.g., Enallopsammia |
|  | Pedal disc | Surface opposite the mouth allowing anchoring to rock bottom through production of adhesive materials. E.g., Anthomasthus, Actiniaria (Clarke et al. 2020). |
|  | Peduncle | Lower part of the Pennatuloidea, a stalk-like structure, embedded in the sediment (Bayer, 1983), or modified peduncle with sucker-shaped over rocky bottom (Williams & Alderslade, 2011). |
|  | Rhizoids | Root-like features extending from the base of the colony (Bayer, 1983) (e.g., Acanella). |
| Skeleton flexibility | Fleshy | No calcified skeleton (e.g., Actiniaria). |
|  | Semi-fleshy | No calcified skeleton but rigidifying parts (e.g., some Zoantharia). |
|  | Semi-rigid | Rigid axis, and the remaining parts are more flexible (e.g., some Zoantharia, Pennatuloidea, Antipatharia, Alcyonacea). On images one can observe some bent parts by the effect of the currents. |
|  | Rigid | Almost completely calcified skeleton (e.g., Stylasteridae, Scleractinia), thick rigid/woody appearance on images. |
| Polyp size | Small | Not visible on the images. |
|  | Medium | Quite visible on the images (one can guess a small cap or a star shape – i.e., polyp – (e.g., Epizoanthus) |
|  | Large | Discernible on the images with clearly visible tentacles (e.g., Anthomastus) |
| Body size | Small | 2-5 cm |
|  | Medium to large | >5-15 cm |
|  | Very large | >>15 cm |
| Spacing between branches | Branching compact | Branches in clusters or tufts, dense appearance, little space among branches (adapted from Veron, 2000). |
|  | Branching opened | Branches of similar length given off at similar angles, open space among branches (adapted from Veron, 2000), "holes" clearly visible among the branches. E.g., some Plexauridae. |
|  | No branch | e.g., single autozoid polyp such as Bathyalcyon, seapen leaf (Pennatuloidea). |
| Number of axes | Single | A single central inner rigidified supporting structure, bearing lateral branches, leaf or polyps. E.g., Pennatuloidea, Chrysogorgia. |
|  | Multiple | Multiple inner rigidified supporting structures. E.g., Enallopsammia |
|  | No main axis | All forms carried by a pedal disc (e.g., Actiniaria) or solitary scleractinians. |
| Polyp arrangement | From a central axis, or leaf | Polyp arrangement from the central axis directly, or leaf extending from the central axis. |
|  | On lateral branches (secondary axes) | Arrangement of the polyp from the lateral (secondary) branches. |
|  | On a basal support | Arrangement of the polyp on top of a large pedal disc or basal support (calcified disc) (e.g., Actiniaria or solitary scleractinians). |
| Internal space/volume | 0/1 | Presence of a semi-enclosed internal space/cavity (e.g., Acanella, Chrysogorgia, colonising Actiniaria on hermit crab). |
| Bends with the water flow | Erect | Structure facing the current, rather right. |
|  | Curved | Structure observed bent by the flow of water. |
| Fan (Pinnate)<br>(CAAB 11 168913) | 0/1 | A single or multiple central axes with parallel side branches in a single plane (Althaus et al., 2014). Branched colonies in which the branching pattern is feather-like, with the branchlets in one plane (Bayer, 1983). E.g., Bathypathes |
| Quill<br>(CAAB 11 168917) | 0/1 | Single stem with polyp leaves, base bulbous. Stem never branched (Althaus et al., 2014). (Pennatuloidea) |
| Arborescent<br>(e.g., CAAB 11 168904, CAAB 11 168911, CAAB 11 077906) | 0/1 | Tree-like structure, networks of branches with multiple axes (2D or 3D), rigid or flexible, regularly organised or not. E.g., Enallopsammia |
| Bushy<br>(e.g., CAAB 11 168908) | 0/1 | Dense, with many short branches, and compact and fluffy appearance (Althaus et al., 2014). |
| Polyp-AS (anchoring small/not visible)<br>(e.g., CAAB 11 290902, CAAB 11 290903) | 0/1 | Only the polyp is visible, on/near the bottom. (E.g., Actiniaria, Scleractinia) |
| Polyp-AL (anchoring large/visible)<br>(e.g., CAAB 11 229901, CAAB 11 229902, CAAB 11 168910) | 0/1 | Polyp(s) with visible large short pedal disc or calcified stem and disc. (E.g., Actiniaria, Alcyoniidae) |
| Bottle-brush<br>(e.g., CAAB 11 168905) | 0/1 | 3D-column with a bushy appearance, short branches radiating from the main axis (Althaus et al., 2014). (E.g., Chrysogorgia, Acanella) |
| Whip<br>(CAAB 11 168917) | 0/1 | A long thin single stem/axis, sometimes twists around itself, with very short lateral branches around the axis or axial polyps directly (Althaus et al., 2014). |
| Umbel-shaped<br>(e.g. CAAB 11 001000,<br>for solitary hydroid). | 0/1 | A calyx formed by the extension of structures (banching/tentacles) at the end of a long stem/pedal disc. E.g., Umbellula, Bathyalcyon, solitary hydrozoa. |

We integrated additional traits identified in the literature, usually considered as adaptations to deep-sea environments, and/or with respect to their ecological function and/or having potential effect on the community. For sponges, we added the sponge’s size, the osculum structure (from their visibility in the images), the level of sponge compactness (that do not reflect necessarily their robustness, which needs manual manipulation), the presence/absence of internal cavities, and of outgrowths/long apparent spicules (i.e., hispidity). For cnidarians, we considered the modularity, anchoring mode to the substratum, relative polyp(s) size, relative cnidarian body size, qualitative measure of spacing between the branches (compact, opened), bending of the structure with the current, as well as provision of an internal volume for other organisms.

#### Classification of the morphotypes into morpho-functional groups

We listed all the morphotypes annotated on the images in a table, and each morphotype was scored according to the set of traits defined previously (**Tables 1 and 2**, ecological relevance of the traits detailed in the **Supplem. mat. 1 and Supplem. mat. 2**).

We scored 1 or 0 according to the presence or absence of the trait for the morphotype considered, to obtain a binary (presence/absence) matrix of ‘morphotypes x traits’.

Morpho-functional groups (MFGs) were delimited using Hierarchical Agglomerative Clustering (HAC) with Unweighted Pair Group Method with Arithmetic Mean (UPGMA), using the *hclust()* function (‘stats’ package, R Core Team, 2021). The HAC was applied on the Gower distance of the ‘morphotype x traits’ matrix of sponges and cnidarians respectively, calculated from the *daisy()* function, using the ‘vegan’ package (Oksanen et al., 2020). We selected the UPGMA clustering method after testing different methods – Ward, Complete, and Simple linkage, Weighted Pair Group Method with Arithmetic Mean, Weighted Pair Group Method with Centroid, Unweighted Pair Group Method with Centroid – based on the best cophenetic correlation value (Borcard et al., 2018).

We selected a cutting threshold of the tree according to the analysis of the silhouette width giving the optimal number of clusters (i.e., measure the degree of membership of a morphotype to its cluster relying on the average dissimilarity between the morphotype and all others belonging to the same cluster) (Borcard et al., 2018). However, we explored other clusters when also having a high average value of silhouette width and giving more ecological coherent partitions. We kept the threshold quite low to retain details in the morphological and functional description of sponges/cnidarians. The choice of the threshold was a trade-off between the description of the morphological diversity and the number of replicates within groups. The numerical classification approach provides a simplified group number from a more complex multi-traits matrix (Tsakalos et al., 2019).

For the sponge morphotypes classification, we removed the ‘sieve plates’ trait from the matrix because of the lack of consistency of this trait annotation (hardly visible on the images).

Then, we exported the ‘images x morphotypes’ abundance matrix of cnidarians and sponges (from BIIGLE annotation platform) and we summed the abundances of morphotypes belonging to the same MFG (i.e., clusters obtained from the HAC). Thus, we obtained a new ‘images x MFGs’ abundance matrix.

From this matrix, we calculated and standardised the MFG densities per polygon of 200 m^2^ (∼60 m linear) sample unit, from the navigation of the camera and its speed, using the ADELIE software. We therefore obtained a final ‘polygons x MFGs’ matrix. In the text and figures, the MFGs of sponges will be referred to as P#, whilst those of cnidarians will be translated to C#.

#### Environmental dataset

Specifics on the environmental data processing and dataset are available in the Material and Methods section in Hanafi-Portier et al. (2024) (summary of the abiotic context of each site in **Supplem. mat. 3**). Current over three depth layers and chlorophyll a concentrations were considered as hydrological parameters. A high resolution bathymetric grid (10 m x 10 m for all sites except for Mayotte island slopes at 20-m resolution) (Audru et al., 2006; SHOM data) was used to calculate seafloor topographical variables: (1) Bottom position index (BPI) at fine (30, 60, 90 m) and large (120, 250, 500 m) scales, (2) longitudinal, transverse and total curvature, (3) aspect, which reflect seabed orientation (i.e., degree of seafloor exposure to water movement) converted to unitless value of eastness and northness (bounded between - 1 and 1) from the cosine and sine of the aspect angle, (4) bottom roughness using an Arc Chord method, and (5) slope. We also classified four geomorphological classes (volcanic, carbonate, sedimentary, mixed) from acoustic (backscatter) data ground-thruthed by images, and quantified substrate composition from manual and semi-automatic image analyses, substrate diversity (from the Shannon-Weaver index) and hardness score (ranging from 1 to 6) (see the list and definitions of variables in **Supplem. mat. 4**). Environmental variables were aggregated into 200 m^2^ sampling units (polygons) using R.

### Statistical analyses

All analyses were performed using the R environment (V4.1.2) (R Core Team, 2021), and all graphical representations were made using the ‘ggplot’ package (Wickham et al., 2016). The data and scripts used to perform the analyses are available at: https://doi.org/10.17882/108256 (Hanafi-Portier et al., 2025).

#### Morpho-functional groups and taxa richness, beta diversity, and composition

We compared MFGs and taxa richness among seamounts and between seamounts and island slopes from sample-based rarefaction curves (on polygons) using the *specaccum()* function with the ‘exact’ method (‘vegan’ package, Oksanen et al., 2020).

We quantified Beta diversity (BD) (here referring to the degree of spatial MFGs or taxa differentiation among sampling units) for each transect using the *beta.div()* function (‘adespatial’ package, Dray et al., 2022) from the Hellinger distance, as well as derived LCBD (Local Contribution to Beta Diversity) and SCBD (Species Contribution to Beta Diversity) indices. The LCBD index reflects the uniqueness (high or low) of sampling units in their composition in MFGs or taxa (unique sites), while the SCBD index reflects the level of variation of each MFG or taxa within the considered transect (MFGs or taxa with high SCBD coefficient have strong contribution in generating variability within transect) (Legendre & De Cáceres, 2013).

#### Morpho-functional groups and taxa responses to environmental constraints, and their roles on megabenthic community structure

We tested the role of environmental drivers in shaping MFGs and taxa composition and spatial structure within and among sites from partial redundancy analysis (pRDA). We used Hellinger-transformed MFG densities, considering different covariables: latitude, longitude, current, and chlorophyll a concentration, to quantify their contributions in isolation. We first removed collinear variables using Pearson pairwise correlations (removal when Pearson r > 0.85) since RDA analysis is sensitive to collinearities between variables. Partial RDA were performed using the *rda()* function and we tested the significance of the pRDA models, axes and environmental variables with *anova.cca()* by a permutation test of the F-statistic (999 permutations, p < 0.05). We selected parsimonious models from forward selections of the explanatory variables with the *ordiR2step()* function (all the previous functions with the ’vegan’ package), using the adjusted R^2^ selection criterion (Borcard et al., 2018).

We quantified the contribution of the environmental variables and their interrelationships from variation partitioning analyses using *varpart()* function (‘vegan’ package) for hydrology (current, Chla), topography and substrate variables, and geomorphology, on the sponge and cnidarian MFGs as well as taxa. We also quantified the contribution of the sponge and cnidarian MFGs or taxa, environmental and spatial variables, and their interrelationships on the megabenthic community structure. We tested the fractions explained with a permutation test (*anova.cca()* function, n = 999 permutations, p-value < 0.05) and represented them as Venn diagrams.

We also assessed the correlation between sponge and cnidarian MFGs and taxa with the megafaunal community, here by considering sponge and cnidarian MFGs or taxa as explanatory variables, accounting for the effect of the environmental drivers and spatial variables (as conditional variables) using pRDA.

## Results

### Description of the morpho-functional groups

From the hierarchical agglomerative clustering (HAC) applied on the 84 sponge and the 84 cnidarian morphotypes that we manually classified from images, we obtained 37 sponge and 34 cnidarian MFGs (**Table 3**, **Table 4**). The high number of MFGs obtained results from the low threshold chosen for the HAC, and we obtained 35% (cnidarians) to 43 % (sponges) of MFGs composed of singleton (only one morphotype), that corresponded to sponges and cnidarians with particular morpho-trait composition, either identified at high taxonomic resolution (e.g., *Bathyalcyon*, Stylasteridae, *Acanella*), or at low resolution (e.g., Porifera-P2 and P5, Demospongiae-P17; Actiniaria-C7, Alcyonacea-C25) (**Table 3**, **Table 4**). The sponge MFG P35 (*Aphrocallistes*) gathered three morphotypes, and the cnidarian MFG C28 (*Umbellula*) gathered two morphotypes, but only different by their size (from small to large forms). For cnidarians, which could be identified at finer taxonomic resolution than sponges, some traits corresponded to taxa belonging to specific taxonomic groups (e.g., quill morphology and peduncled anchoring mode for the Pennatuloidea although it included nine morphotypes within two MFGs - C33, C34) (**Table 2**). Other MFGs gathered different taxa (C3, C11, C12), of which C13 included taxa all belonging to the Alcyonacea order (Alcyonacea, Primnoidae, Holaxonia, Isididae) and an unidentified colonial anthozoan.

**Table 3.**
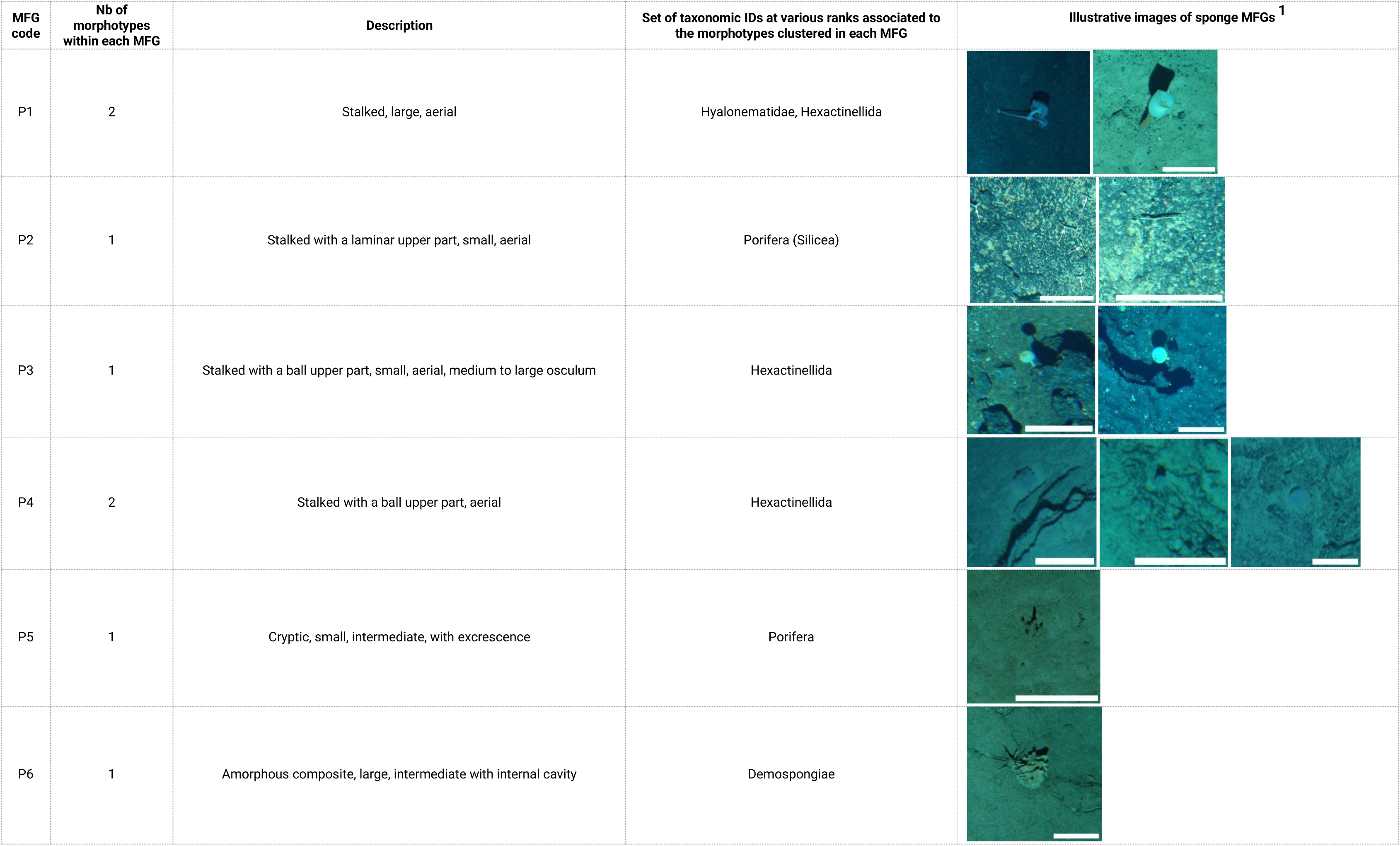
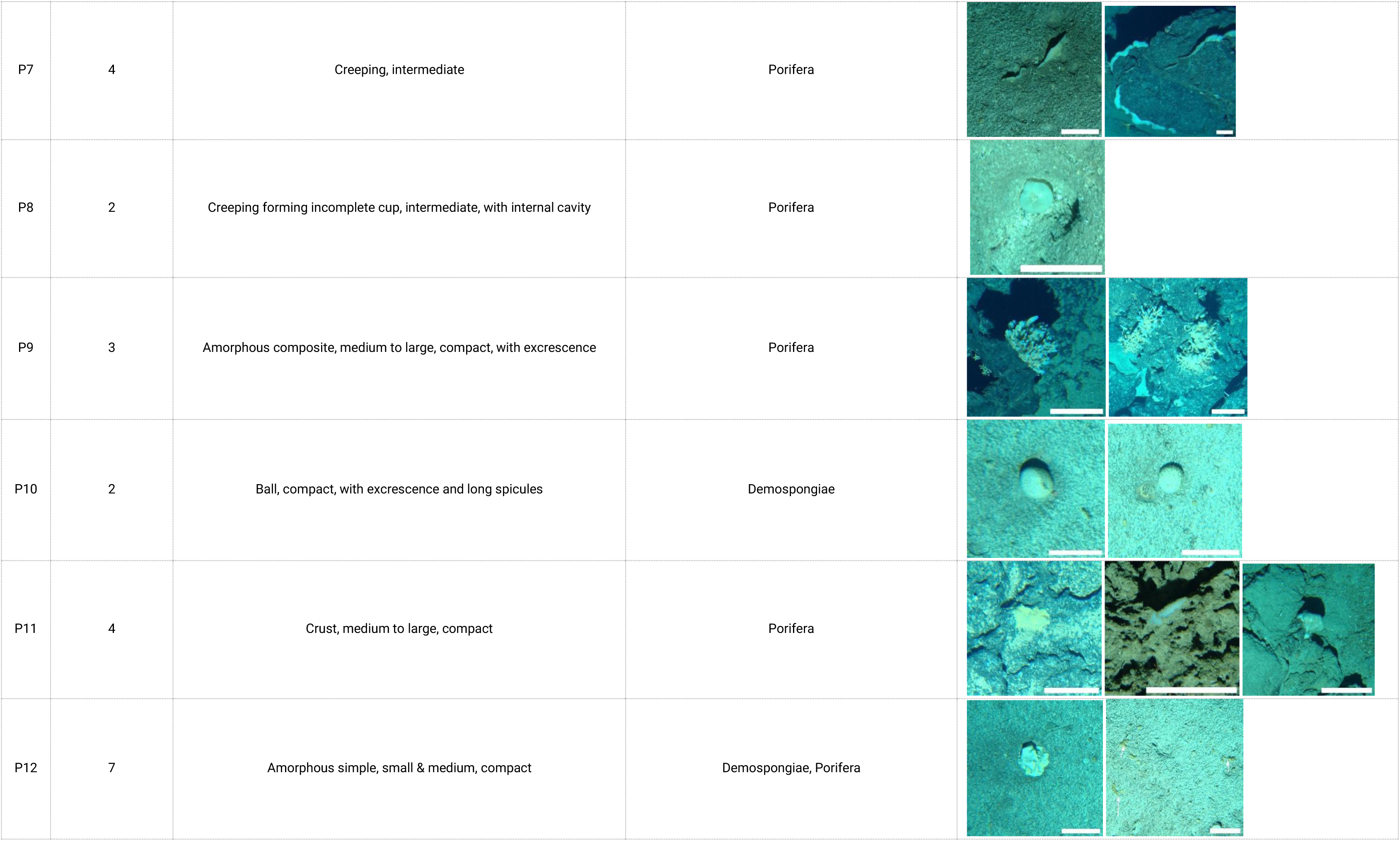
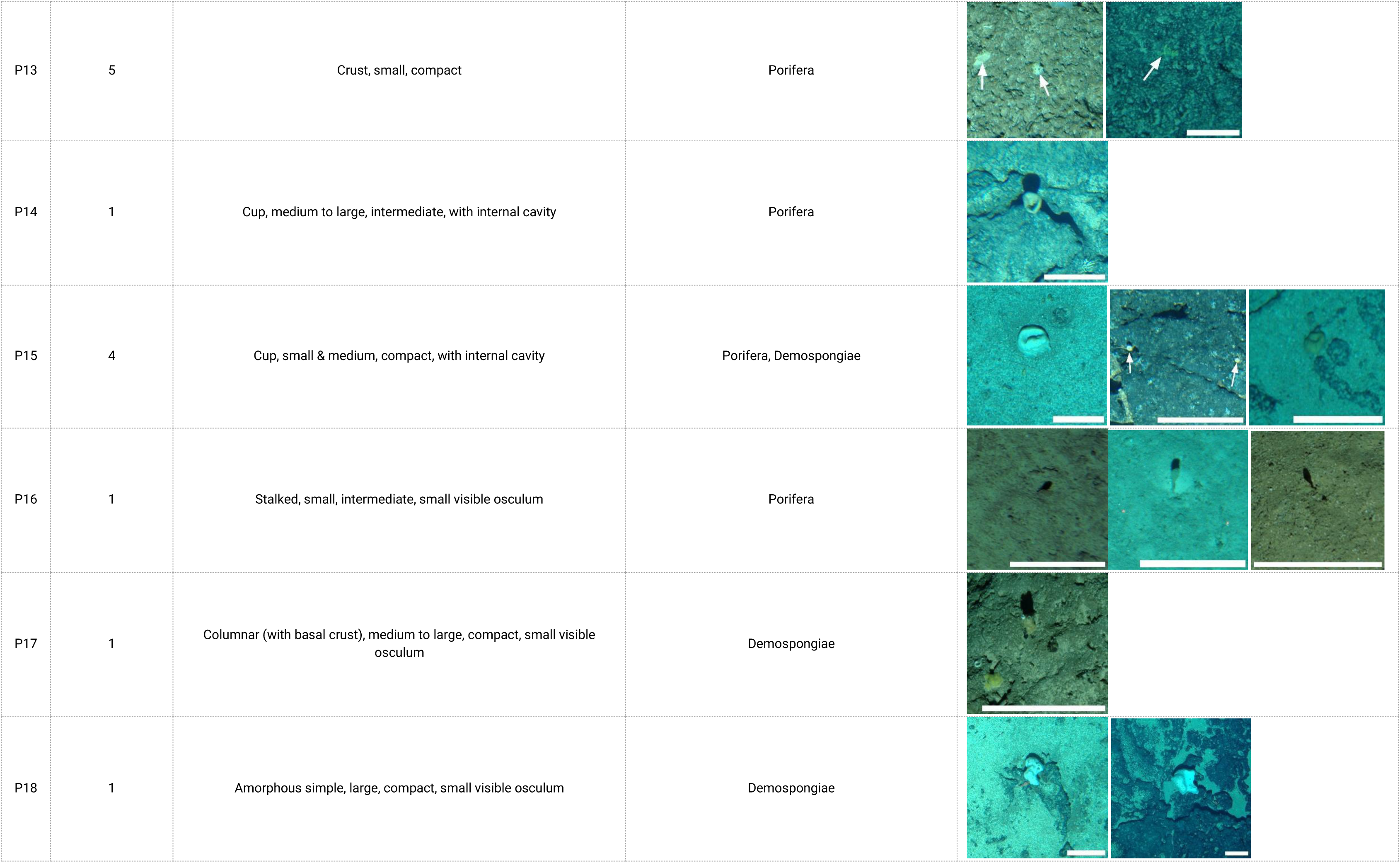
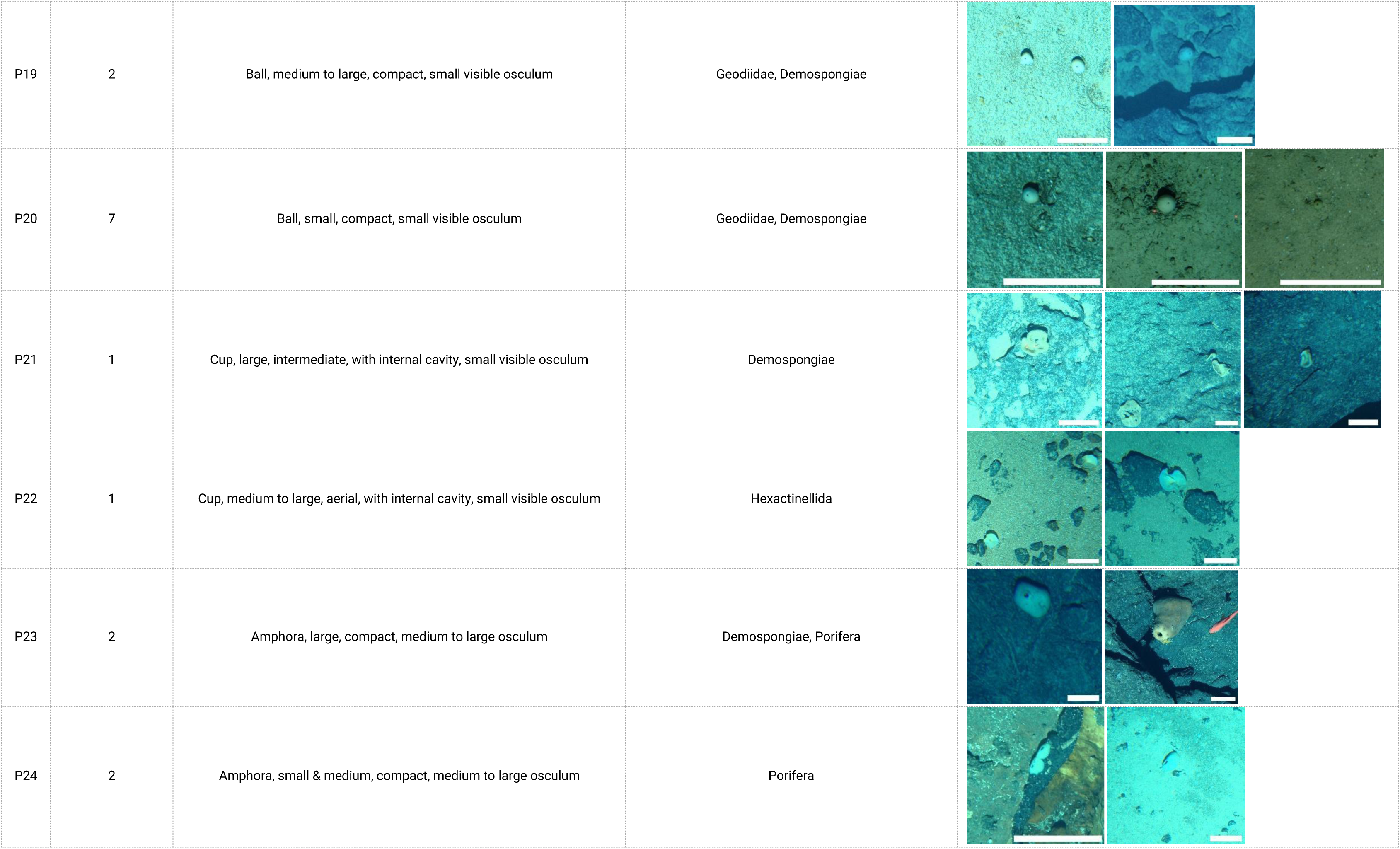
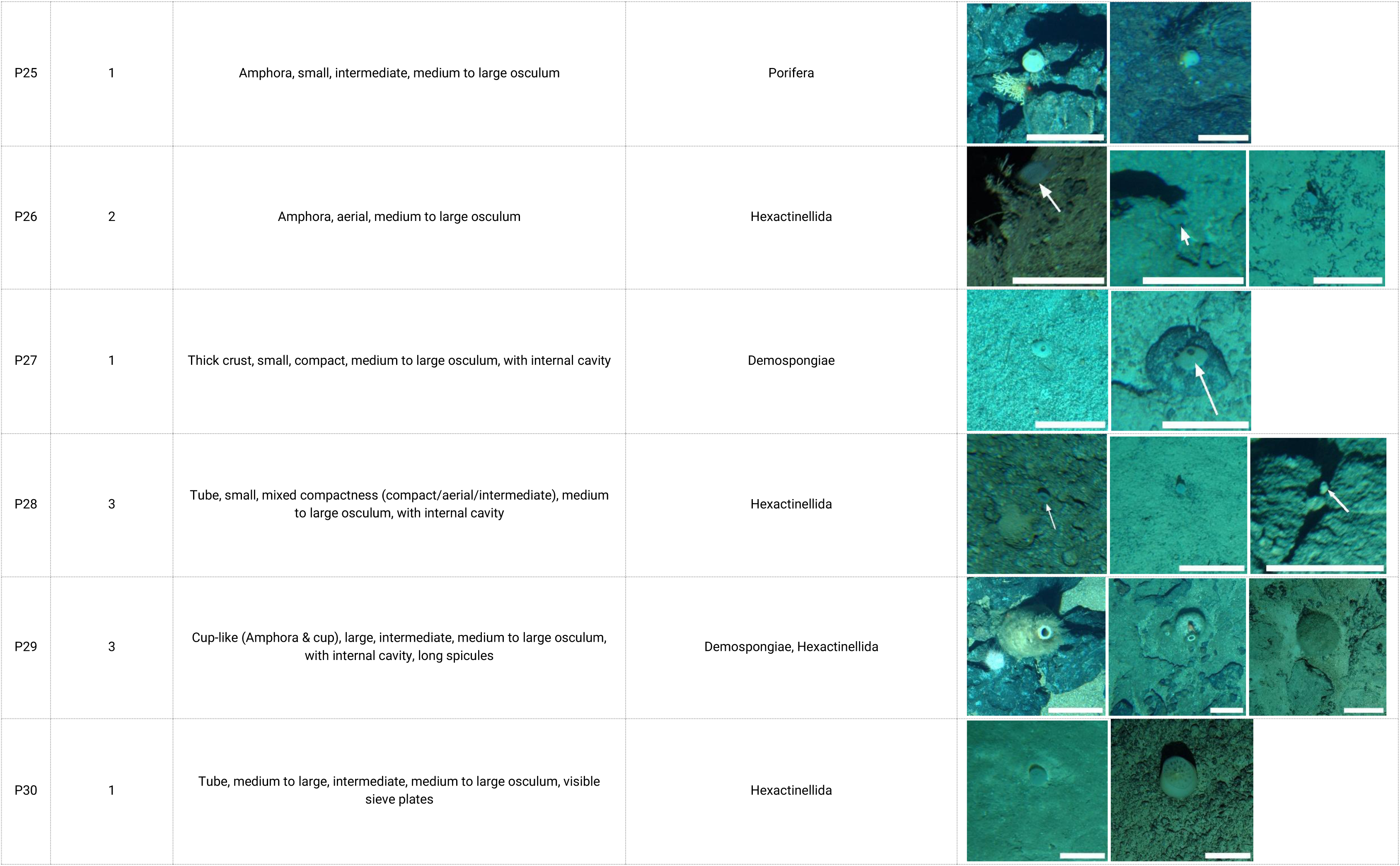
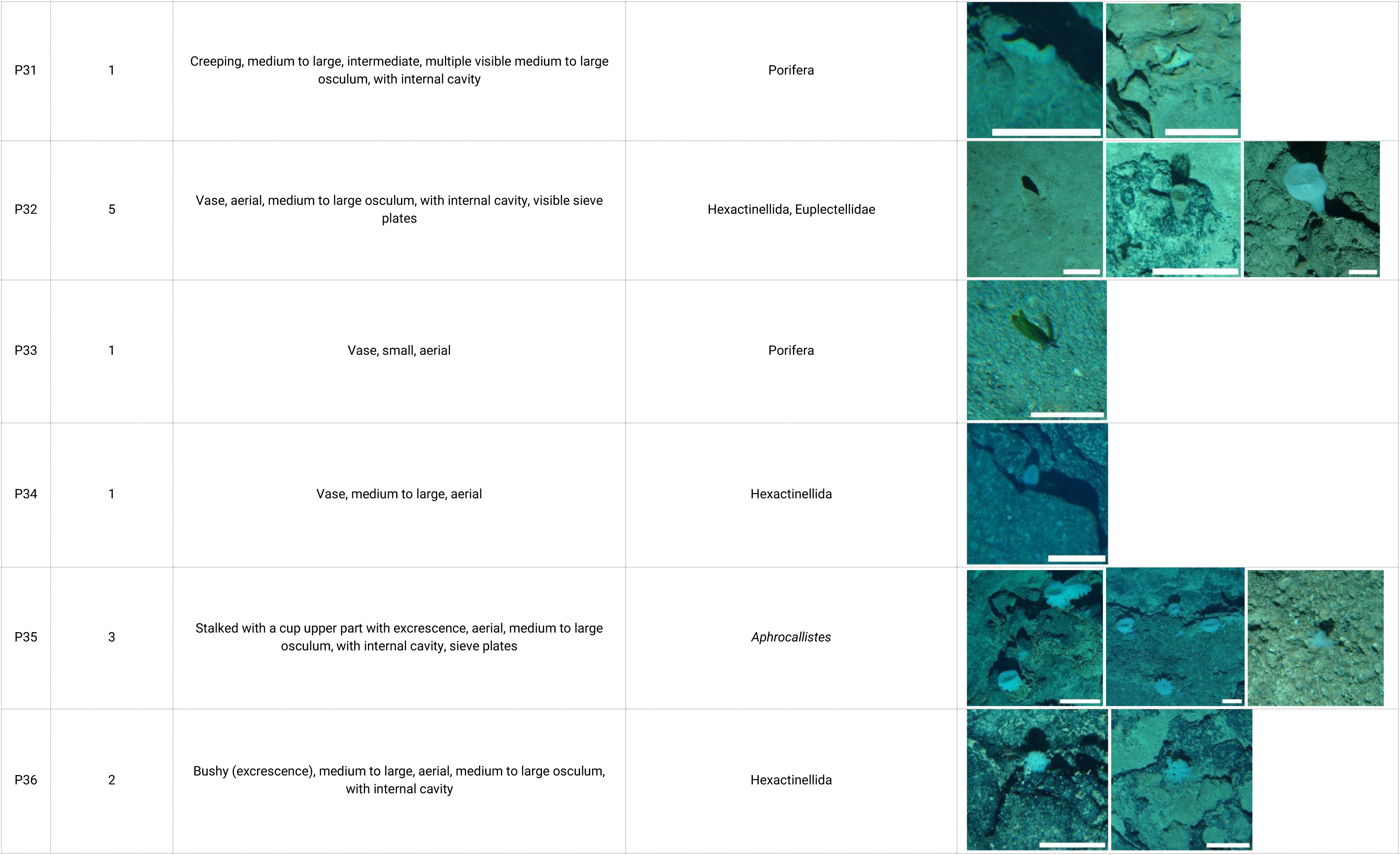
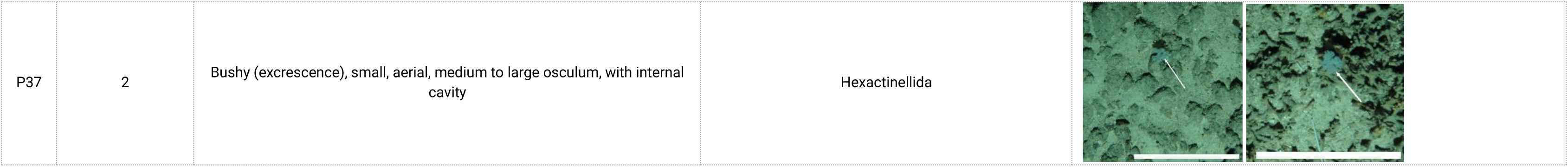
Sponge morpho-functional groups obtained from hierarchical agglomerative clustering.

| MFG code | Nb of morphotypes within each MFG | Description | Set of taxonomic IDs at various ranks associated to the morphotypes clustered in each MFG | Illustrative images of sponge MFGs <sup>1</sup> |
| --- | --- | --- | --- | --- |
| P1 | 2 | Stalked, large, aerial | Hyalonematidae, Hexactinellida |  |
| P2 | 1 | Stalked with a laminar upper part, small, aerial | Porifera (Silicea) |  |
| P3 | 1 | Stalked with a ball upper part, small, aerial, medium to large osculum | Hexactinellida |  |
| P4 | 2 | Stalked with a ball upper part, aerial | Hexactinellida |  |
| P5 | 1 | Cryptic, small, intermediate, with excrescence | Porifera |  |
| P6 | 1 | Amorphous composite, large, intermediate with internal cavity | Demospongiae |  |
| P7 | 4 | Creeping, intermediate | Porifera |  |
| P8 | 2 | Creeping forming incomplete cup, intermediate, with internal cavity | Porifera |  |
| P9 | 3 | Amorphous composite, medium to large, compact, with excrescence | Porifera |  |
| P10 | 2 | Ball, compact, with excrescence and long spicules | Demospongiae |  |
| P11 | 4 | Crust, medium to large, compact | Porifera |  |
| P12 | 7 | Amorphous simple, small & medium, compact | Demospongiae, Porifera |  |

|  |  |  |  |
| --- | --- | --- | --- |
| P13 | 5 | Crust, small, compact | Porifera |
| P14 | 1 | Cup, medium to large, intermediate, with internal cavity | Porifera |
| P15 | 4 | Cup, small & medium, compact, with internal cavity | Porifera, Demospongiae |
| P16 | 1 | Stalked, small, intermediate, small visible osculum | Porifera |
| P17 | 1 | Columnar (with basal crust), medium to large, compact, small visible osculum | Demospongiae |
| P18 | 1 | Amorphous simple, large, compact, small visible osculum | Demospongiae |
| P19 | 2 | Ball, medium to large, compact, small visible osculum | Geodiidae, Demospongiae |
| P20 | 7 | Ball, small, compact, small visible osculum | Geodiidae, Demospongiae |
| P21 | 1 | Cup, large, intermediate, with internal cavity, small visible osculum | Demospongiae |
| P22 | 1 | Cup, medium to large, aerial, with internal cavity, small visible osculum | Hexactinellida |
| P23 | 2 | Amphora, large, compact, medium to large osculum | Demospongiae, Porifera |
| P24 | 2 | Amphora, small & medium, compact, medium to large osculum | Porifera |
| P25 | 1 | Amphora, small, intermediate, medium to large osculum | Porifera |
| P26 | 2 | Amphora, aerial, medium to large osculum | Hexactinellida |
| P27 | 1 | Thick crust, small, compact, medium to large osculum, with internal cavity | Demospongiae |
| P28 | 3 | Tube, small, mixed compactness (compact/aerial/intermediate), medium to large osculum, with internal cavity | Hexactinellida |
| P29 | 3 | Cup-like (Amphora & cup), large, intermediate, medium to large osculum, with internal cavity, long spicules | Demospongiae, Hexactinellida |
| P30 | 1 | Tube, medium to large, intermediate, medium to large osculum, visible sieve plates | Hexactinellida |

|  |  |  |  |
| --- | --- | --- | --- |
| P31 | 1 | Creeping, medium to large, intermediate, multiple visible medium to large osculum, with internal cavity | Porifera |
| P32 | 5 | Vase, aerial, medium to large osculum, with internal cavity, visible sieve plates | Hexactinellida, Euplectellidae |
| P33 | 1 | Vase, small, aerial | Porifera |
| P34 | 1 | Vase, medium to large, aerial | Hexactinellida |
| P35 | 3 | Stalked with a cup upper part with excrescence, aerial, medium to large osculum, with internal cavity, sieve plates | *Aphrocallistes* |
| P36 | 2 | Bushy (excrescence), medium to large, aerial, medium to large osculum, with internal cavity | Hexactinellida |

|  |  |  |  |
| --- | --- | --- | --- |
| P37 | 2 | Bushy (excrecence), small, aerial, medium to large osculum, with internal cavity | Hexactinellida |

**Table 4.**
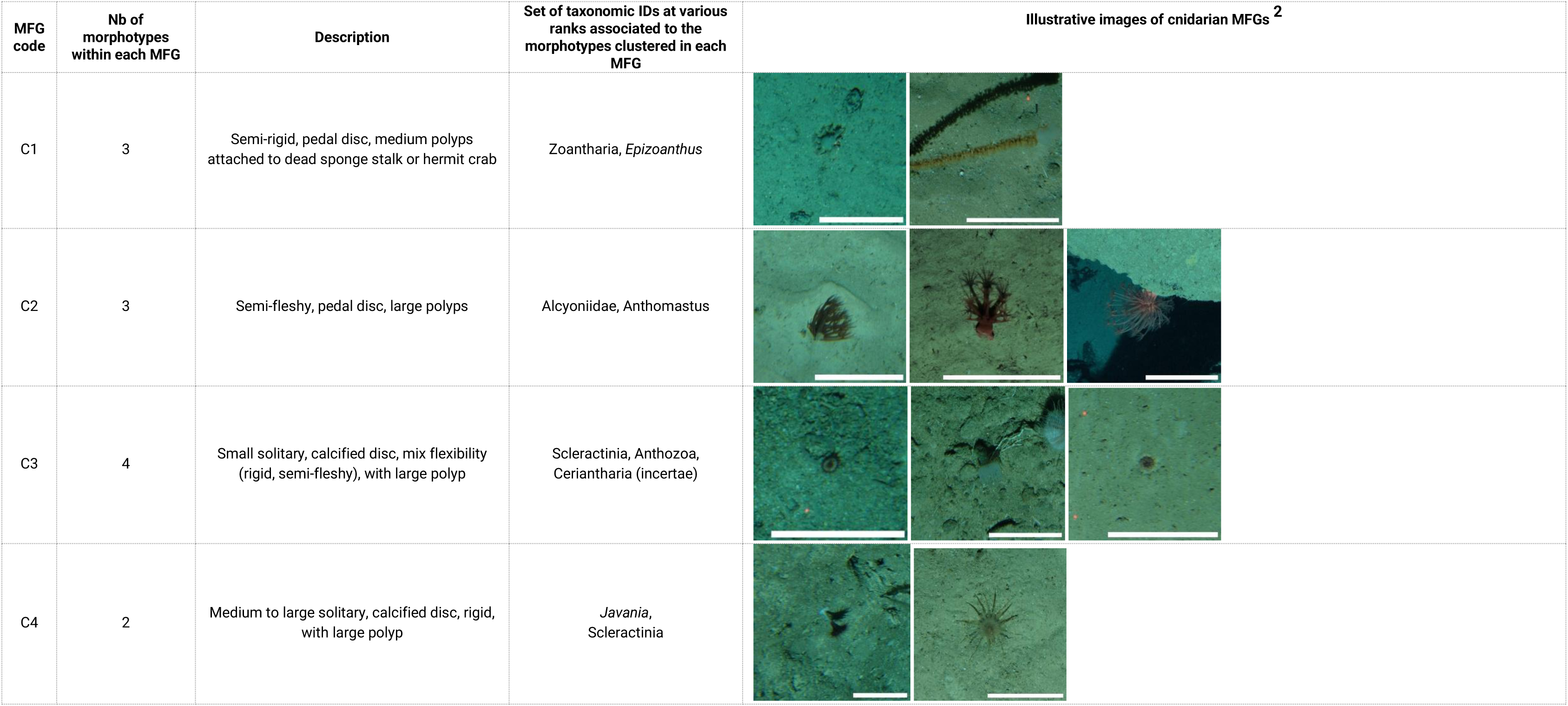
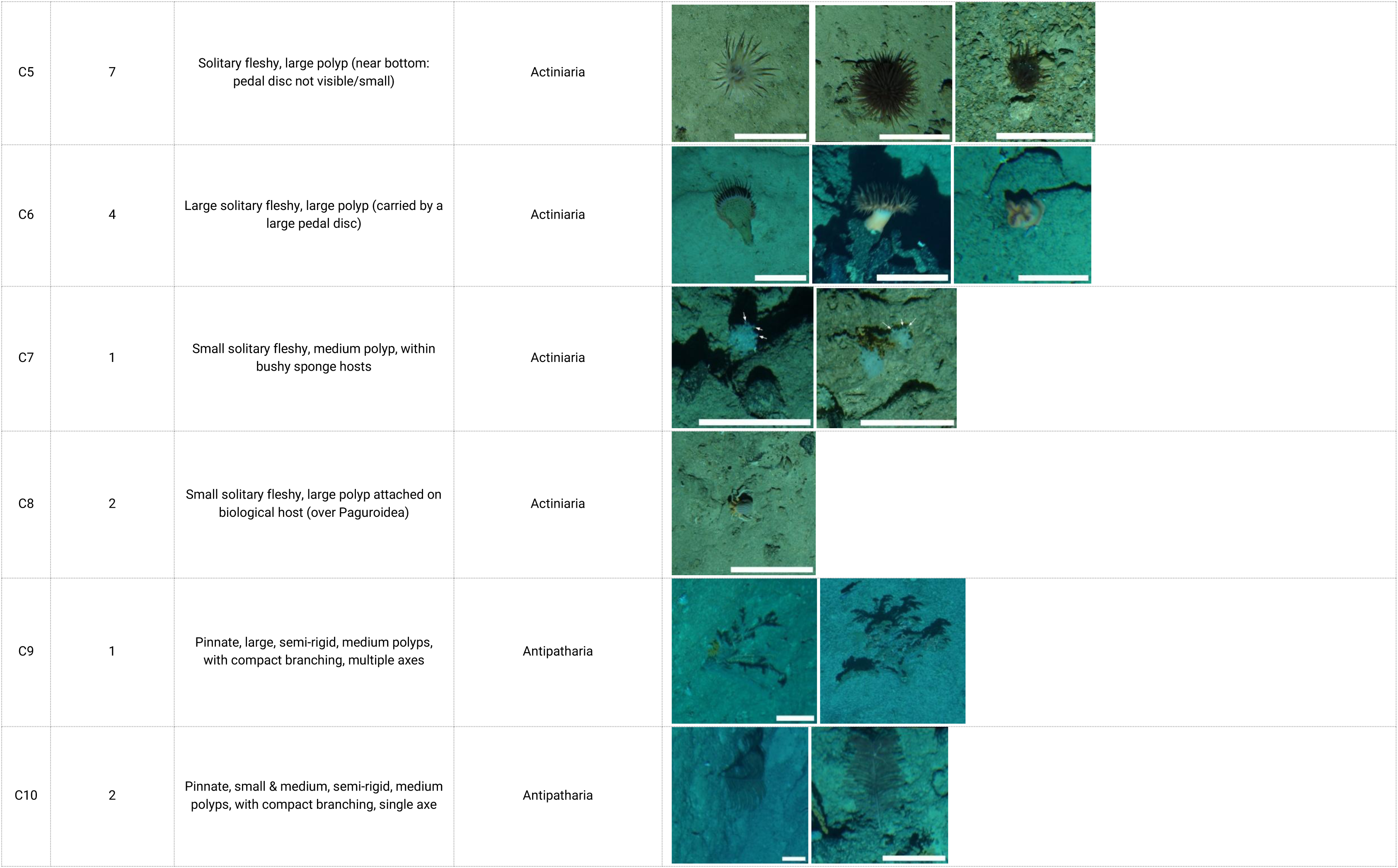
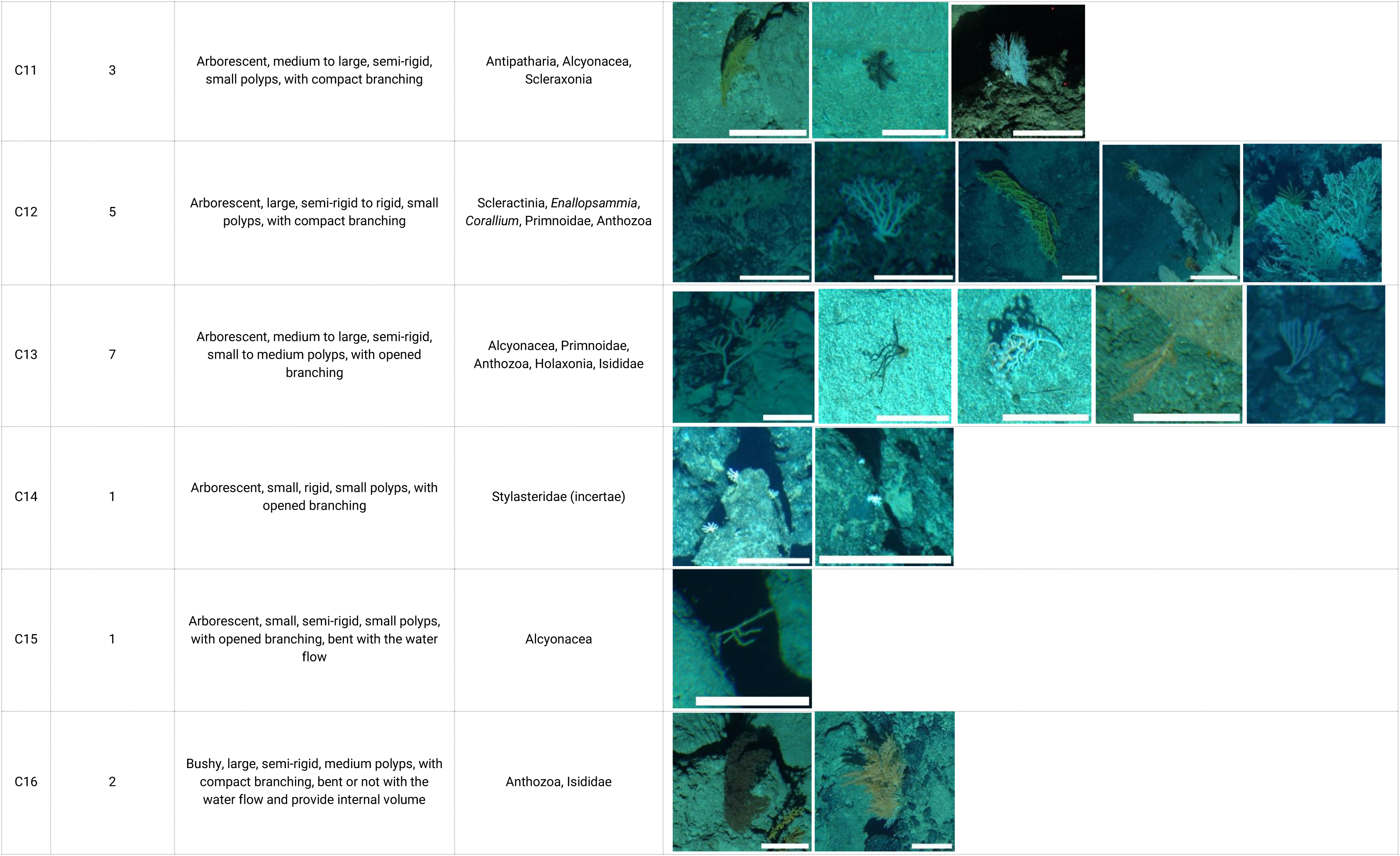
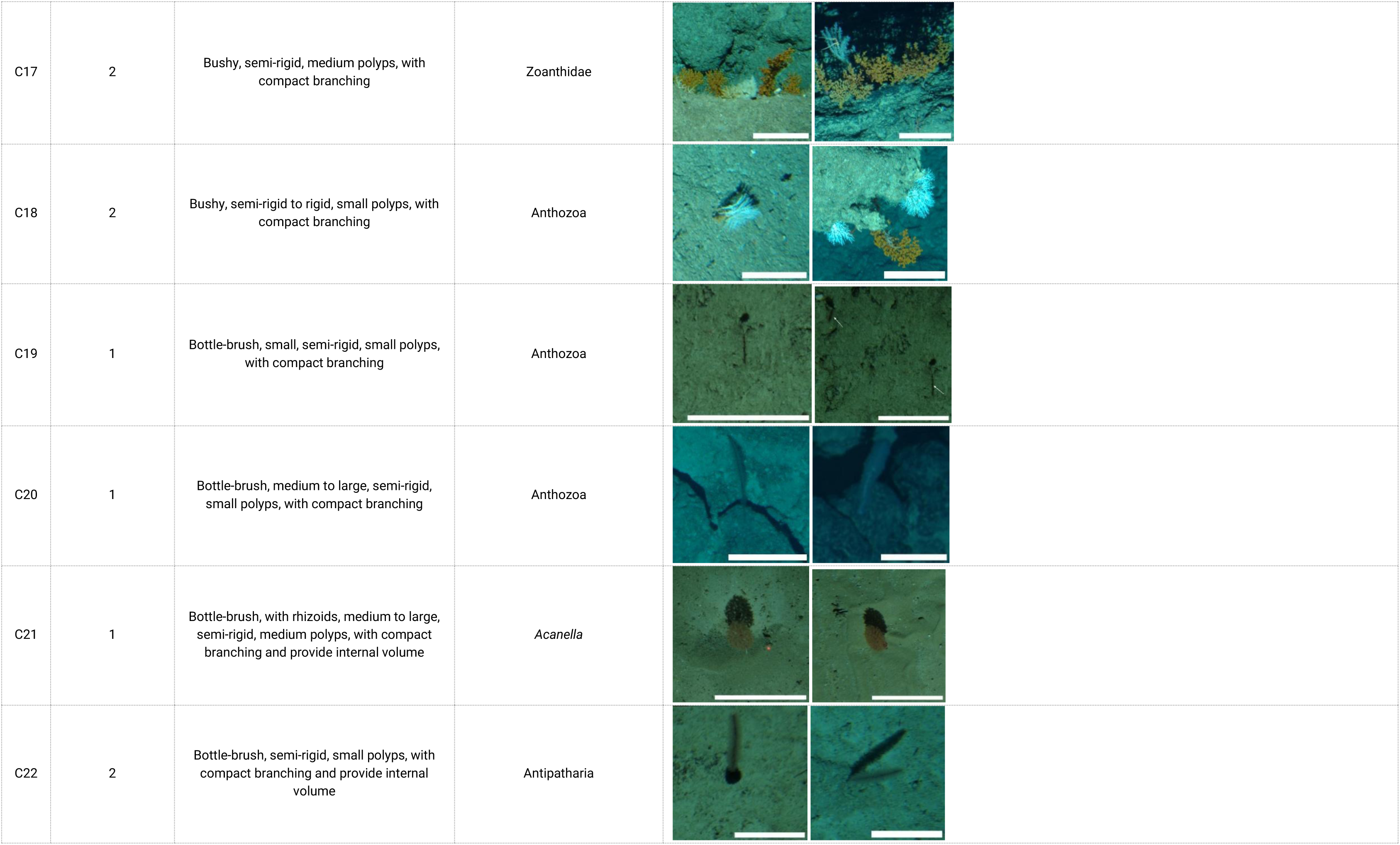
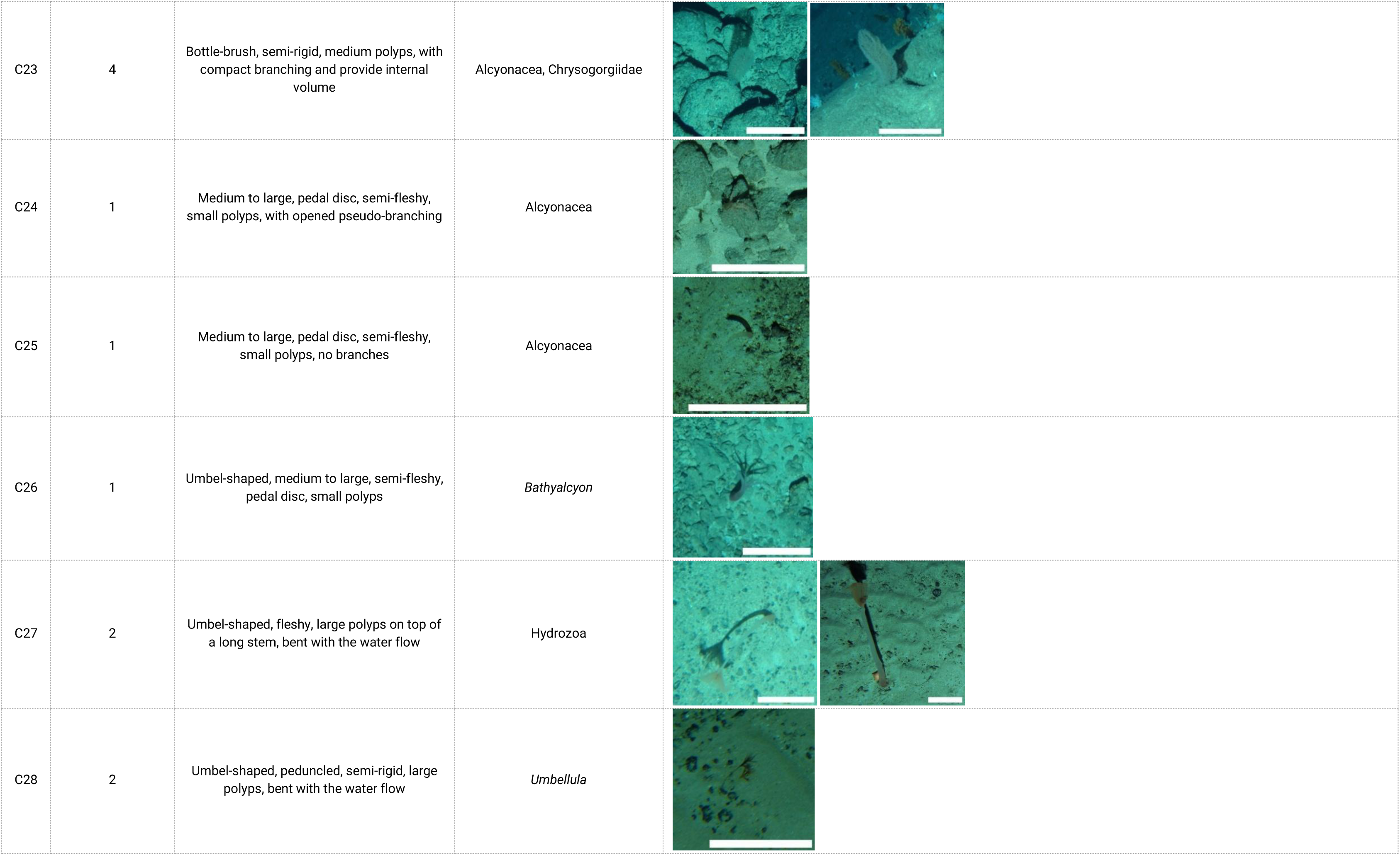
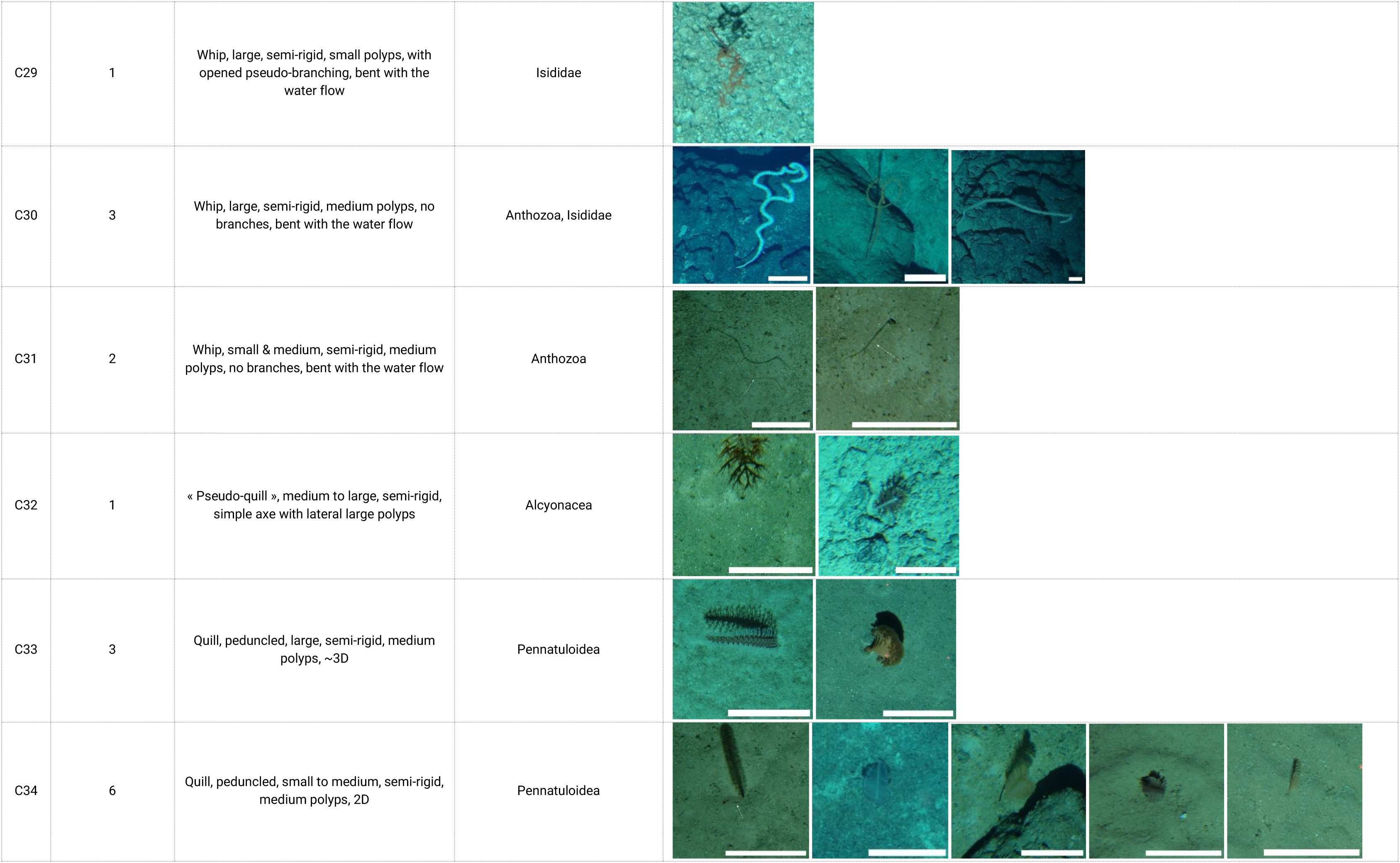
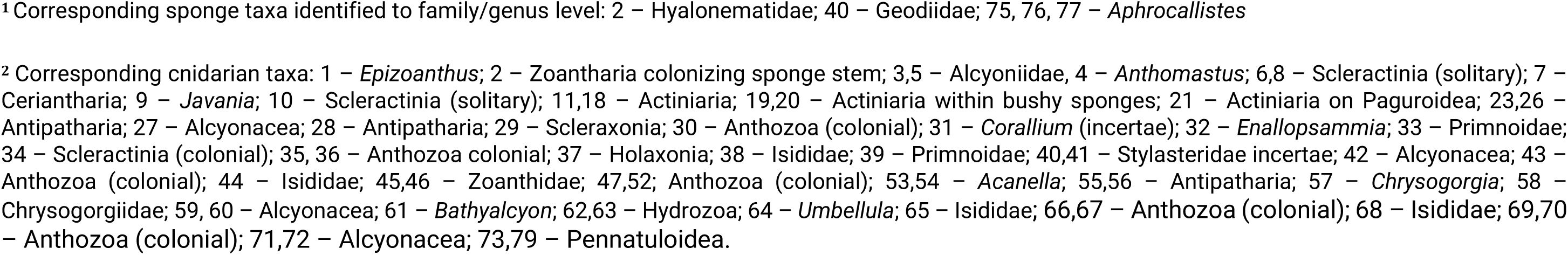
Cnidarian morpho-functional groups obtained from hierarchical agglomerative clustering.

### Quantitative comparison of the Morpho-functional *vs.* Taxonomic Classifications

To objectively evaluate the relevance of an MFG framework compared to coarse-resolution taxonomic classification schemes, we compared richness and beta diversity metrics (Table 5).

**Table 5.**
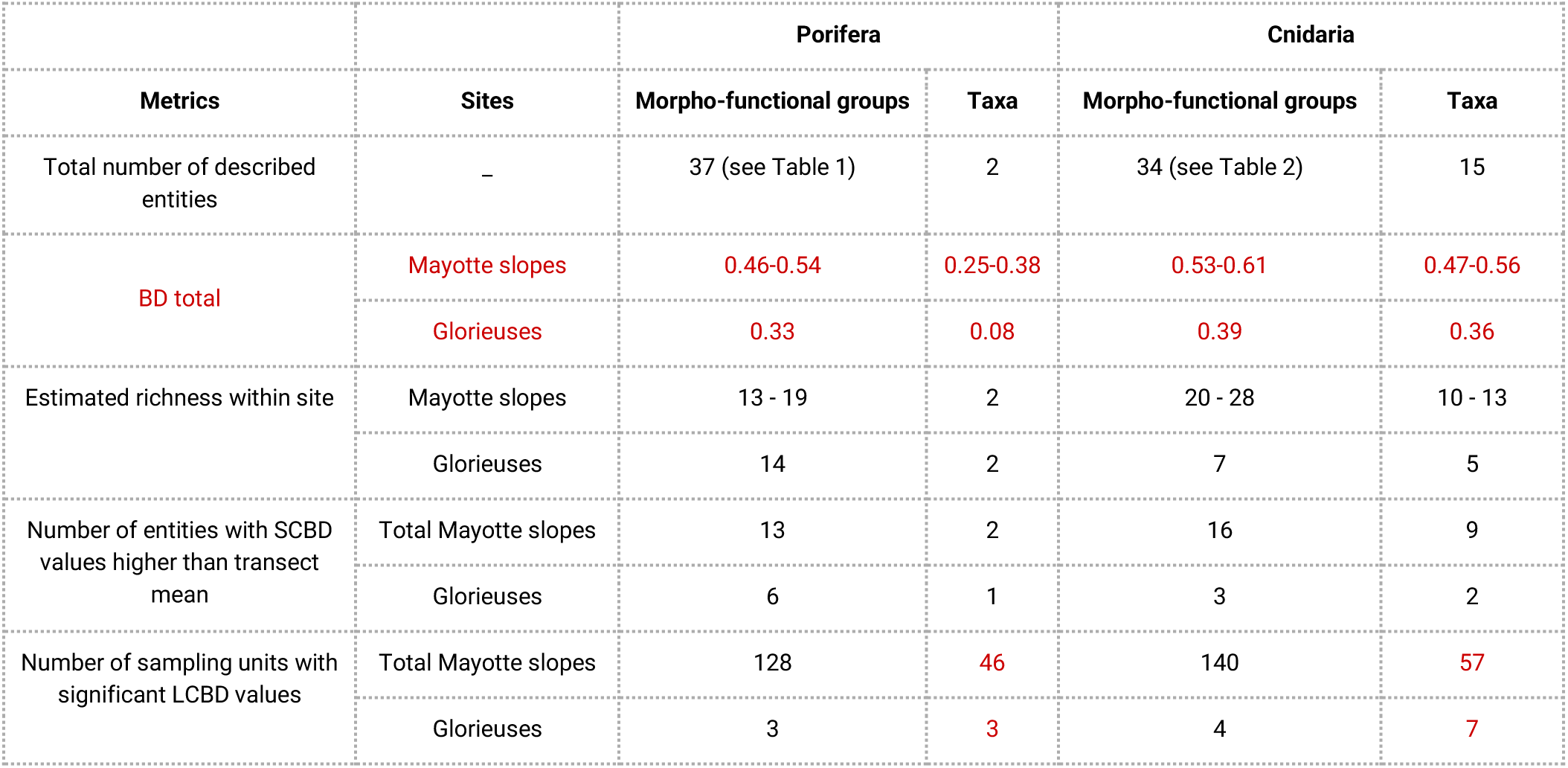
Comparison of richness (cnidarians: n = 106; sponges: n = 58 polygons), total Beta Diversity, SCBD* and LCBD** of sponges and cnidarians, between the morpho-functional and the taxonomic classifications with examples for sites in the northern Mozambique Channel. *’Species Contribution to Beta Diversity’ as the degree of variation of a species across the study area. **’Local Contribution to Beta Diversity’ as the uniqueness of the site.

| Metrics | Sites | Porifera |  | Cnidaria |  |
| --- | --- | --- | --- | --- | --- |
|  |  | Morpho-functional groups | Taxa | Morpho-functional groups | Taxa |
| Total number of described entities | – | 37 (see Table 1) | 2 | 34 (see Table 2) | 15 |
| BD total | Mayotte slopes | 0.46-0.54 | 0.25-0.38 | 0.53-0.61 | 0.47-0.56 |
|  | Glorieuses | 0.33 | 0.08 | 0.39 | 0.36 |
| Estimated richness within site | Mayotte slopes | 13 - 19 | 2 | 20 - 28 | 10 - 13 |
|  | Glorieuses | 14 | 2 | 7 | 5 |
| Number of entities with SCBD values higher than transect mean | Total Mayotte slopes | 13 | 2 | 16 | 9 |
|  | Glorieuses | 6 | 1 | 3 | 2 |
| Number of sampling units with significant LCBD values | Total Mayotte slopes | 128 | 46 | 140 | 57 |
|  | Glorieuses | 3 | 3 | 4 | 7 |

### (A) Diversity patterns and spatial distribution of sponge and cnidarian morpho-functional groups *vs.* taxa

#### MFG and taxa composition

The composition and relative frequencies of sponge and cnidarian MFGs varied among surveyed sites (**Figures 2**, **Supplem. mat. 5A, 5B**). There were about eight main sponge morphologies – dominant at one or several sites – which differed in occurrence and relative frequencies among sites: P36, P12, P37, P2, P15, P21, P22, P35. (**Figure 2A**). The MFG approach for sponges allowed for better differentiation of sites, as there were three to ten main (most frequent) morphologies per site, compared to two classes.

**Figure 2:**
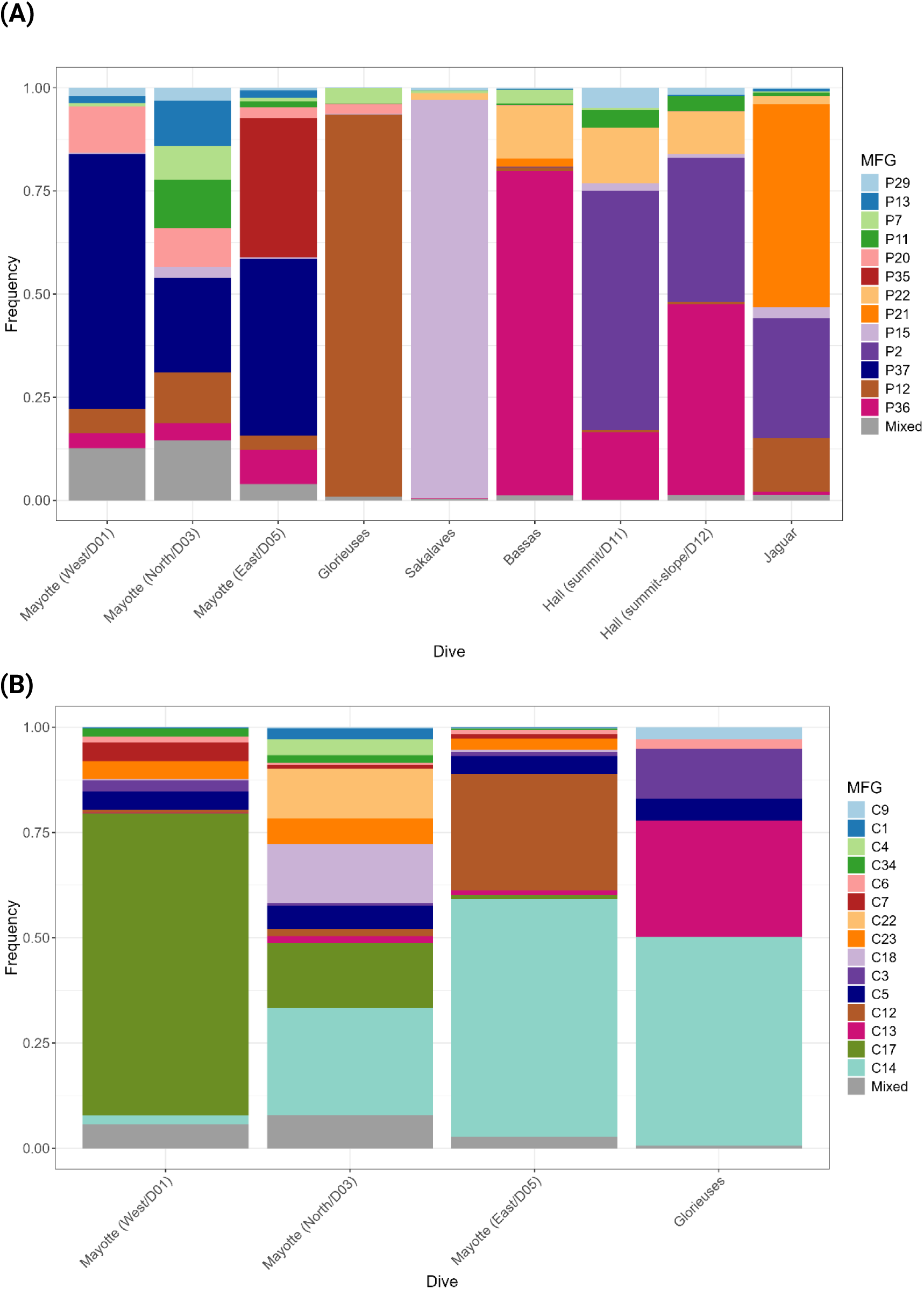
Histograms of the (A) sponge and (B) cnidarian MFGs representing 95% of the total frequency among sites. In the legend, the top labels represent the least frequent MFGs while at the base the most frequent MFGs. Apart is the “mixed” category that regroups MFGs representing the remaining 5% of total frequency (rare MFGs).

For cnidarians in the northern area, we observed five abundant morphologies along Mayotte island slopes and on Glorieuses, which differed in relative frequencies among sites: C14, C13, C5 and C3 (on all sites) (**Figure 2C**). Other morphologies were restricted to Mayotte island slopes, such as C17, C12, C18, C23, or C7, but differences in relative frequencies were observed among the three slopes. The composition of sites with the most frequent dominant MFGs roughly corresponds to the composition of cnidarian orders (**Supplem. mat. 6A, 6B**).

#### MFG and taxa richness

At equal sampling effort (n = 58 polygons), the richness of sponge and cnidarian MFGs differed among sites (**Supplem. mat. 7**).

The highest sponge MFGs richness were observed on the northern slope of Mayotte (19 MFGs) and the eastern slope (16 MFGs) while the lowest were observed on Sakalaves, Bassas da India, and Hall summit (7 to 8 MFGs) (**Supplem. mat. 7A**). Richness of sponge classes (taxa) was not surprisingly equal of two for all sites along the Mozambique Channel (**Table 5**, **Supplem. mat. 7C**),

For cnidarians, at equal sampling effort in the northern area (n = 106 polygons), the highest MFGs richness was observed on the northern slope of Mayotte (28 MFGs) as for sponges, while the richness was very low on Glorieuses (7 MFGs) (**Supplem. mat. 7B**). The cnidarian taxa richness was higher and similar on Mayotte island slopes (10-13) compared to Glorieuses (5 taxa) (**Table 5, Supplem. mat. 7D**). Patterns of richness differences among sites did not differ between the two classifications (cnidarian MFGs or taxa).

#### Variability of morpho-functional group and taxa assemblages within and among sites

The spatial variability of MFGs differed among sites (beta diversity).

For sponges, the MFG beta diversities (BD) were the highest along the eastern slope of Mayotte and the summit-slope of Hall Bank (0.54 and 0.51 respectively), while they were the lowest along Glorieuses and Sakalaves platforms and Bassas external slope (0.33 to 0.39). Similarly, the sponge taxa BD were the highest along the eastern slope of Mayotte (0.38) and the summit and summit-slope of Hall Bank (0.33-0.35), and the lowest along Sakalaves platforms and Glorieuses (0.08-0.13). The range of total beta diversity of sponge MFGs was almost two to four-fold higher than for sponge taxa (**Table 5**).

For cnidarians in the northern Mozambique Channel, the MFG BDs were the highest along the eastern and northern slopes of Mayotte (0.60-0.61 out of 1) and the lowest along the Glorieuses transect (0.39). The beta diversity of cnidarian taxa ranged similarly than with cnidarian MFGs, and with the same order of magnitude (from 0.36 along Glorieuses to 0.56 along the northern slope of Mayotte) (**Table 5**).

The number of sponge and cnidarian MFGs with high SCBD coefficient differed among sites (i.e., MFGs generating more variability than the transect average) (**Supplem. mat. 8**). Some sites had many sponge and cnidarian MFGs with high contribution to the BD (e.g., Mayotte island slopes) while others had a few (e.g., Sakalaves and Bassas for sponges, Glorieuses for cnidarians).

Comparison with SCBD values calculated at the taxonomic level (**Table 5**, **Supplem. mat. 8**) showed that for sponges, only one taxon was detected as having high SCBD coefficient at each site, consistent with the low taxonomic resolution of this group in our dataset (only two classes). For corals, the MFGs identified as having high SCBD were generally associated with taxa that also displayed high SCBD values, indicating a correspondence between the contribution of MFGs and that of their constituent taxa to beta-diversity.

Similarly, polygons with significant high LCBDs (i.e., unique in their MFG assemblage compositions and which contribute the most to the BD) varied in number and in spatial distribution among sites (**Figure 3A, 3C**). They were numerous along the Mayotte island slopes, distributed over small spatial scales, particularly abundant in volcanic areas, but also on other predominantly carbonate bottoms and locally in sedimentary areas (where blocks of hard substrate were sparsely distributed). Conversely, on the Glorieuses, and for sponges also along the seamounts of the central/southern Mozambique Channel, we identified less polygons with significant LCBDs, more or less localised in specific areas such as volcanic areas (Sakalaves, Bassas, Hall), in transition areas between geomorphological facies (all sites), sloping areas (Glorieuses) and locally sediment-dominated areas (**Figure 3A, 3C**).

**Figure 3.**
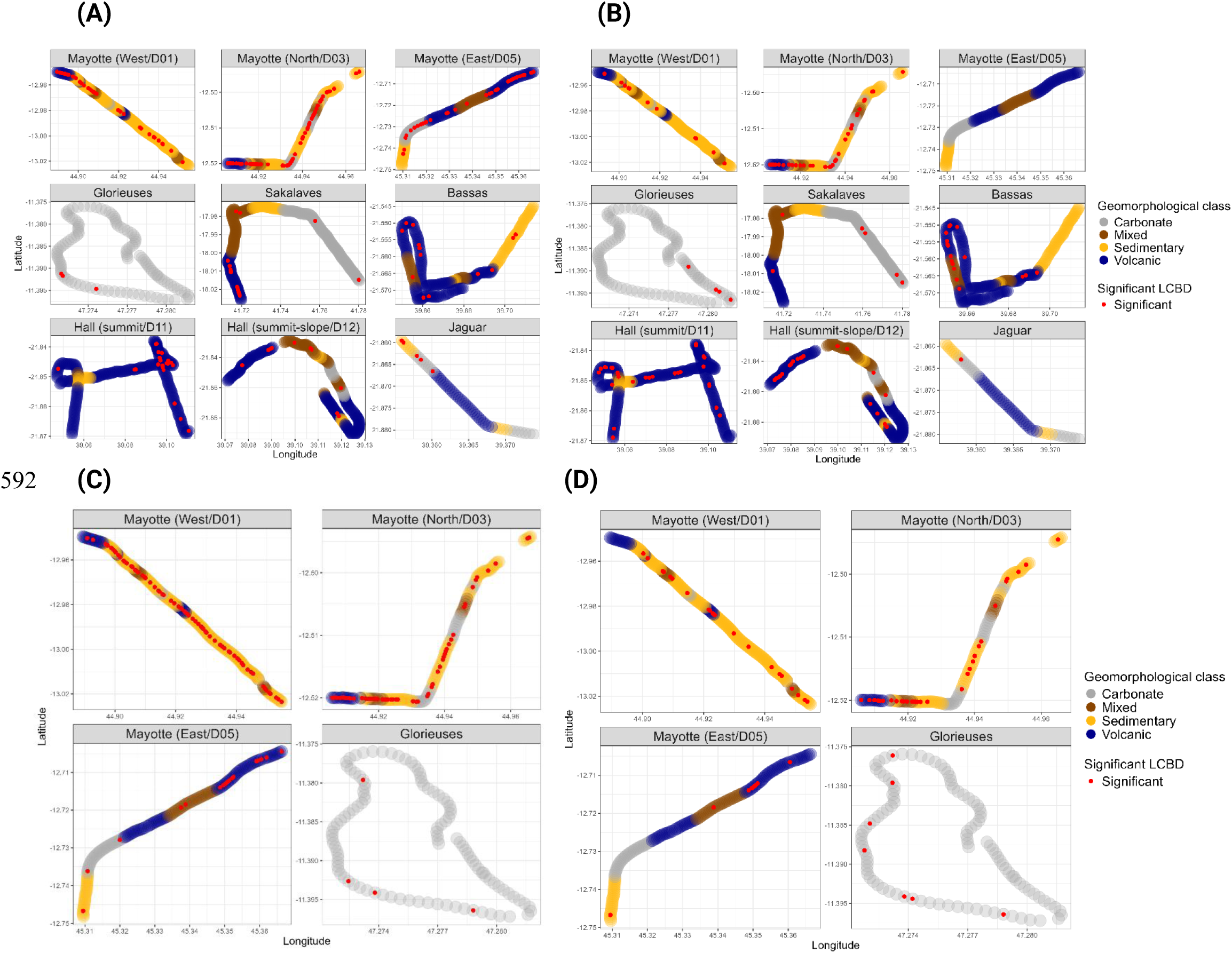
Maps representing, (A) and (B) for sponge MFGs and taxa respectively; (C) and (D) for cnidarian MFGs and taxa respectively. Polygons in red dot having a significant LCBD value (Local Contribution to Beta Diversity, from 999 permutations, p < 0.05) i.e., with a unique MFGs composition, for each transect.

The number of polygons with significant LCBDs was consistently higher when based on sponge and cnidarian MFGs than when based on taxa (**Table 5**). This difference was particularly evident along the eastern slope of Mayotte where no significant LCBD was detected for sponge taxa (**Figure 3B**), whereas numerous significant LCBDs were detected using MFGs (**Figure 3A**). This pattern was also particularly pronounced for both sponges and cnidarians along the Mayotte island slopes, highlighting the greater spatial differentiation in assemblage composition captured by the MFG approach.

### (B) Environmental drivers of sponge and cnidarian morpho-functional groups *vs*. taxa

Sponge MFG composition along the Mozambique Channel was significantly correlated with 19 of the 28 environmental variables tested, which jointly explained approximately 20% of its spatial structure, compared with 17% explained by the spatial variables (latitude and longitude) in the partial RDA (pRDA; **Figure 4A**). Hydrological variables, particularly current variability and inter-annual variability in chlorophyll a, were the main contributors to sponge MFG variance (26.4%), followed by topographic and substrate variables (22.2%; **Supplem. mat. 9A**). Current variability, slope, bathymetry, carbonate geomorphology and substrate hardness were among the variables most strongly associated with MFG composition. Several MFGs showed distinct correlations with environmental conditions. For example, P36 (medium/large bushy aerial) was found in deeper and steeper volcanic areas, and to a smaller extent with higher rugosity, BPIs 120, 500 m, and low deep current speed and variability. P15 (small/medium cup compact) was positively correlated to high current variability, hard and carbonate seabed; P12 (small/medium amorphous simple compact) and P2 (small stalked form with upper laminar part) were positively correlated to strong deep current variability and speed to a less extent (**Figure 4A**). The northern-channel pRDA further discriminated between compact MFGs (e.g., P12, P13, P11, P7) positively correlated with carbonated, hard seabed and higher winter chlorophyll a, and aerial or stalked MFGs (e.g., P37, P36, P35, P32; i.e., small/large bushy aerial, stalked aerial with excrescences, medium/large vase aerial) positively correlated with deeper, sloping volcanic areas (**Supplem. mat. 10A**). The pRDA based on sponge taxa showed a similar balance between environmental and spatial components, with 21.5% and 20.2% of the variance explained, respectively (**Figure 4B**). However, fewer environmental variables were significant than for MFGs, and these were predominantly broad-scale hydrological and environmental descriptors, including winter chlorophyll a concentration, its inter-annual variability, bathymetry, deep current variability and BPI500. A similar pattern was observed in the northern Mozambique Channel, where fewer variables were retained and mainly reflected broad-scale hydrological and geomorphological gradients. In both taxonomic pRDAs, only the first RDA axis was significant and primarily separated the two sponge classes, Hexactinellida and Demospongiae. Variation partitioning further showed a lower proportion of residual variance for the taxonomic approach, both across the Mozambique Channel and in the northern area, with hydrological and geomorphological components contributing more strongly than in the MFG-based models (**Supplem. mat. 9B, 9D**).

**Figure 4.**
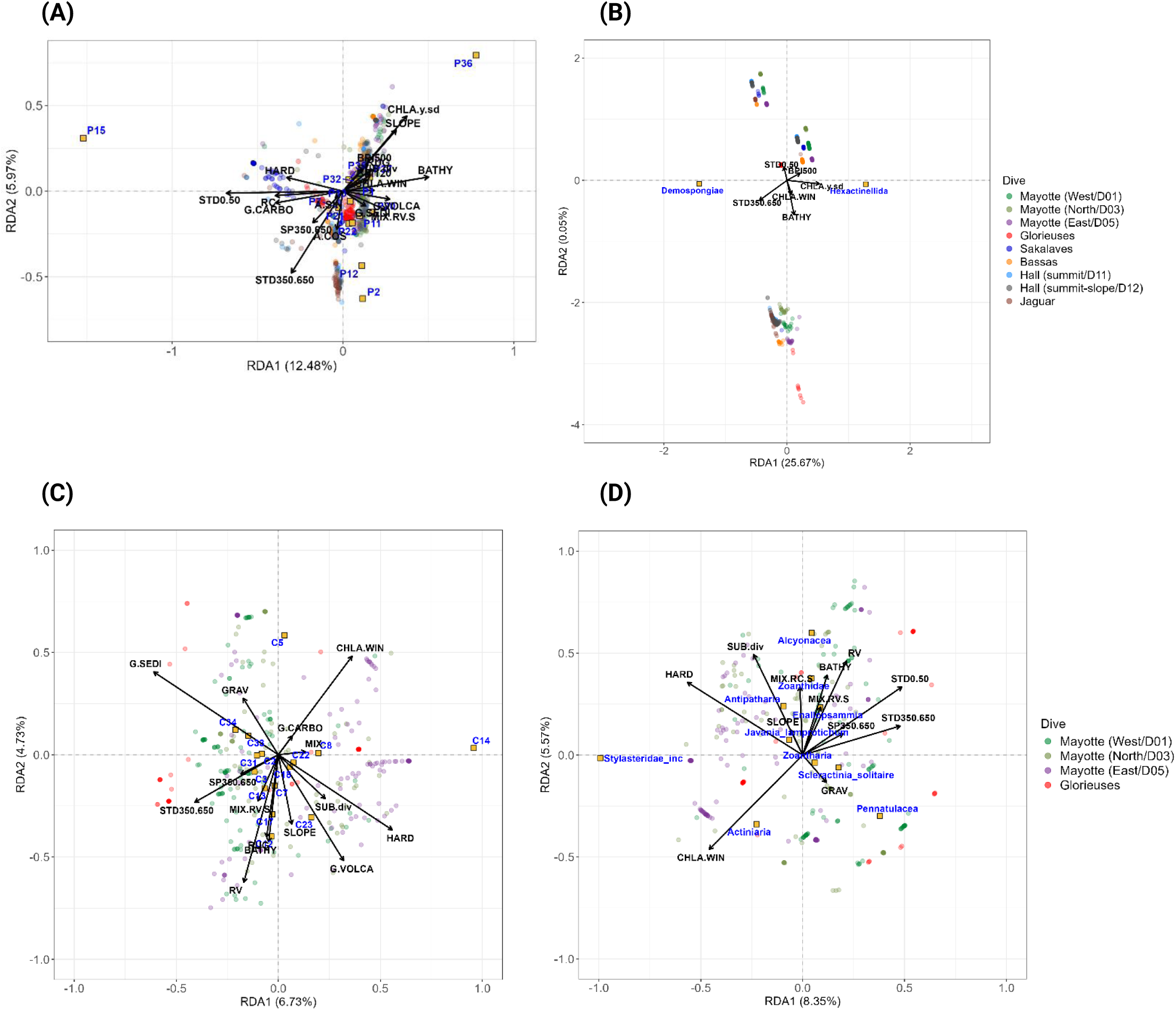
Partial Redundancy Analysis (pRDA) applied to the relative densities of (A) sponge MFG and (B) sponge taxa (classe), along the Mozambique Channel, (C) cnidarian MFGs and (D) cnidarian taxa along the northern Mozambique Channel, after quadratic and Hellinger transformation of the data. The conditional co-variables of the model are latitude and longitude. The significant variables of the model, obtained after a forward selection and test of the significant variables (n = 999 permutations), are shown.

Cnidarian MFG composition was significantly correlated with 15 of the 28 environmental variables tested, which jointly explained approximately 19% of its spatial structure, compared with ∼6% explained by the spatial variables (latitude and longitude) in the pRDA (**Figure 4C**). Sedimentary and volcanic geomorphology, substrate hardness, volcanic rock cover, winter chlorophyll *a* concentration and bottom-current variability were among the strongest environmental correlates. In the northern Mozambique Channel, topographic and substrate variables were the main contributors to MFG variance (∼17%), followed by hydrological variables (7.9%; **Supplem. mat. 9E**). The pRDA mainly highlighted habitat preferences (substrate, geomorphology, topography, current) rather than a clear separation among northern sites. Quill (i.e., C33, 34 - Pennatuloidea) and solitary fleshy MFGs (i.e., C5 - Actiniaria) were positively correlated with gravelly sedimentary habitats, whereas large arborescent, bushy and bottle-brush forms (e.g., C12, C17, C23) were positively correlated with deeper, rugged, sloping hard and volcanic seabed. Some MFGs also showed contrasting associations with current variability, with bushy and large semi-rigid arborescent forms (e.g., C13, C17, C12) associated with higher current variability, while small rigid arborescent (C14 - Stylasteridae (incertae)) and solitary fleshy (C5 - Actiniaria) forms were associated with lower variability and higher winter chlorophyll a concentration (**Figure 4C**).

The pRDA based on cnidarian taxa showed a close correspondence with the MFG-based analysis (**Figure 4D**). Environmental and spatial variables explained similar proportions of community variation (∼21.8% and ∼7%, respectively), and both RDA axes were significant. Twelve environmental variables were significant, with substrate diversity and hardness, volcanic rock cover, bathymetry, winter chlorophyll a concentration and bottom-current variability among the strongest contributors. Variation partitioning likewise showed a similar distribution of explained and residual variance between the MFG- and taxon-based analyses (**Supplem. mat. 9F**), indicating a correspondence between the two classifications for cnidarians.

### (C) Structuring role of sponge and cnidarian morpho-functional groups *vs*. taxa, on megabenthic communities

Variation partitioning (**Figure 5**) was used to disentangle the pure contribution of sponge and cnidarian MFGs or taxa to benthic assemblage variance from the fractions shared with spatial and environmental variables.

**Figure 5:**
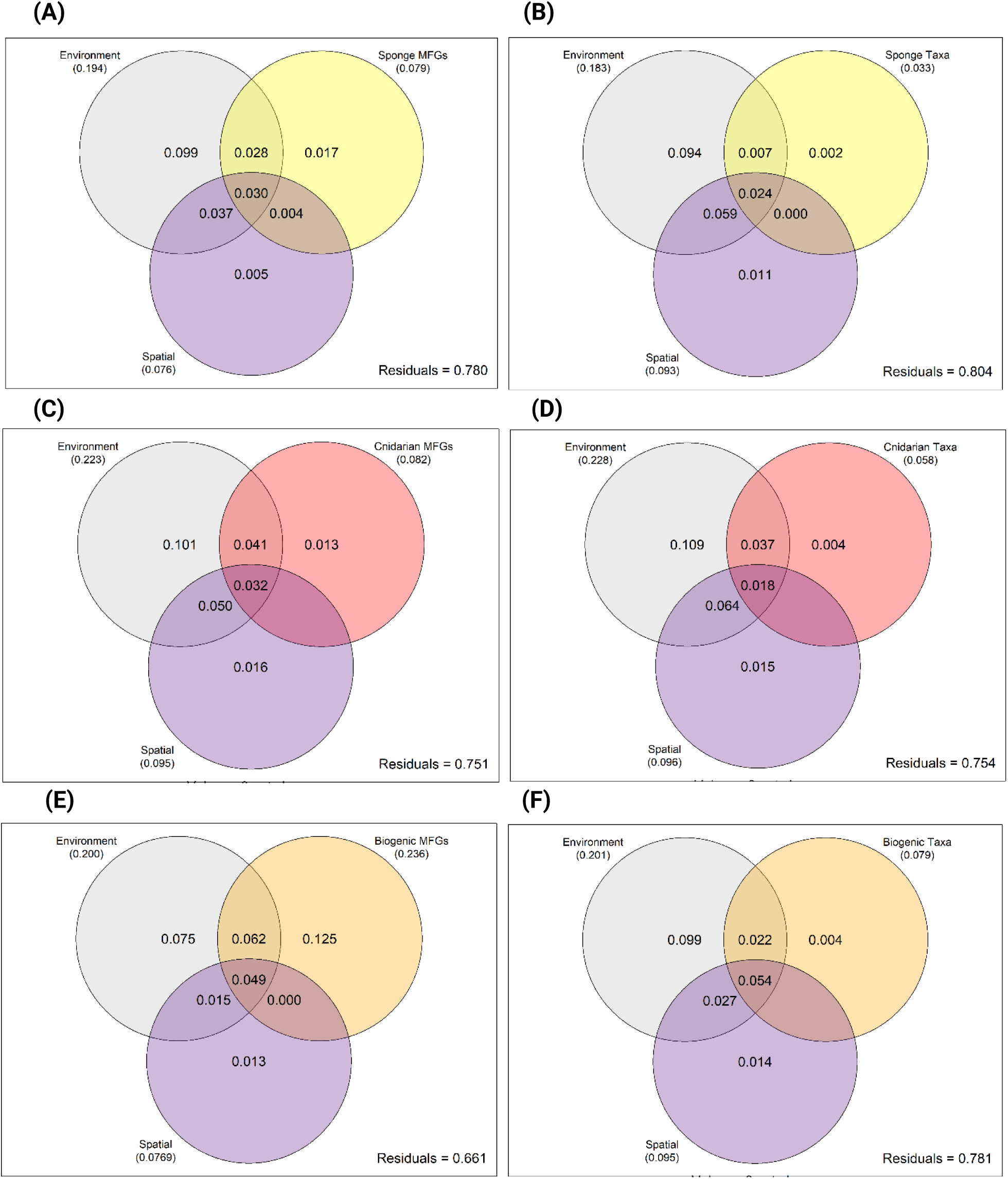
Venn diagrams representing the contributions from the environmental, spatial, and biogenic component to the megabenthic assemblage variance, resulting from a variation partitioning analysis: along the Mozambique Channel for (A) sponge MFGs and (B) sponge taxa; along the northern Mozambique Channel for (C) cnidarian MFGs and (D) cnidarian taxa, and for (E) sponge and cnidarian MFGs and (F) sponge and cnidarian taxa. Numbers indicate the fractions of variation explained by each set of variables. Contributions < 0 are not shown.

Across the Mozambique Channel, sponge MFGs had a significant pure contribution of 1.7%, while their total contribution reached 7.9% when shared fractions with space and environmental variables were included (**Figure 5A**). Environmental variables accounted for the largest proportion of explained variance, with a pure contribution of ∼10% and a total contribution of 19.4%. Overall, 78% of the benthic variance remained unexplained, and all fractions were significant.

The sponge taxon-based variation partitioning showed the same general pattern, but with a much smaller sponge taxa pure contribution (0.2%) and a lower total contribution (3.3%); all fractions were significant (**Figure 5B**).

In the northern Mozambique Channel, the pure sponge-MFG and sponge-taxa fractions and the fraction shared with spatial structure were not significant, and the environmental component remained the dominant contributor in explaining benthic assemblage variance.

For cnidarians, all fractions were significant when MFGs were considered (**Figure 5C**). Their pure contribution accounted for 1.3% of benthic variance, while their total contribution reached 8.2%. Although environmental variables again explained the largest proportion of variance, a significant fraction was shared among environmental variables, spatial structure and cnidarian MFG composition (3.2%).

In contrast, the pure contribution of cnidarian taxa and the fraction shared with spatial structure were not significant (**Figure 5D**). Their total contribution was lower than that of cnidarian MFGs (5.8%). Again, environmental variables accounted for the largest proportion of explained variance, having a pure contribution of ∼11% and a total contribution of ∼23%.

When sponge and cnidarian MFGs were considered jointly, their total contribution was significant and reached 23.6%, higher than the total contribution of the environment (20%). However, their pure contribution was not significant despite accounting for 12.5% of the variance (**Figure 5E**). In contrast, the total contribution of the two taxonomic groups was only 7.9%, with neither the pure taxonomic fraction (0.4%) nor the fraction shared with spatial structure (<0) being significant (**Figure 5F**).

Overall, environmental variables generally accounted for the largest proportion of explained megabenthic assemblage variance, while MFGs generally accounted for a larger total contribution than taxa, particularly when sponge and cnidarian MFGs were considered jointly.

Where the pure fraction of the corresponding MFGs or taxa was significant in the variation partitioning, partial RDAs were used to test the association between their composition and benthic assemblage structure while conditioning for environmental and spatial variables (**Supplem. mat. 12**).

For sponge MFGs along the Mozambique Channel, the pRDA model was significant, with nine MFGs significantly correlated with benthic assemblage structure (P36, P21, P11, P37, P22, P25, P1, P35 and P15; **Supplem. mat. 12A**). The pRDA based on sponge taxa was also significant, with only Hexactinellida significantly correlated with benthic assemblage structure, and only the first RDA axis was significant (**Supplem. mat. 12B**).

The pRDA for cnidarian MFGs was significant (**Supplem. mat. 12C**) with forward selection identifying four MFGs (C8, C31, C1 and C14) significantly associated with benthic assemblage structure.

Several positive correlations between sponge MFGs and benthic assemblages (including cnidarians) were apparent in the pRDA, including correlations between P25, P37 and P35 (aerial amphora, small aerial bushy and large aerial stalked cup forms, respectively) and Zoanthidae, and between P11 (crust) and P36 (large bushy aerial) and Alcyonacea (**Supplem. mat. 12A**). For cnidarians, C1 (Zoanthiaria, *Epizoanthus*) was correlated with *Hexactinellida*, Paguroidea and *Steredocidaris*, while C8 (Actiniaria) was correlated with Paguroidea, an association also apparent in the imagery (**Supplem. mat. 12B**). The benthic taxa most strongly associated with the selected axes were mostly fishes, hermit crabs, small gastropods and epibenthic taxa.

Overall, the variance explained by the selected axes was very low in all pRDAs, and these associations should therefore be interpreted with caution.

## Discussion

We aimed to assess the potential of a morpho-functional approach to characterise habitat-forming-taxa in poorly or sparsely sampled areas and/or with low taxonomic knowledge, and to determine whether morphological information could provide additional insights into their spatial distribution and ecological associations from their visible morphological diversity from imagery. By relying on morphological information, this framework could be readily integrated into a standard classification system, such as CATAMI, to overcome uncertainties and subjectivity in taxonomic identification, as well as variations in taxonomic nomenclature, while generating morphological data and patterns of structural diversity that are comparable across studies. Our results show that MFGs provide further details on the spatial structure of sponge and cnidarian communities over seamounts and island slopes explored in the Mozambique Channel compared to low resolution taxonomic IDs. The results differed between the two taxa.

### Comparison between the morpho-functional and the taxonomic classifications to describe habitat-forming-taxa

The multi-trait statistical classification defined a large number of sponge and cnidarian MFGs, with 43% of sponge and 35% of cnidarians corresponding to a single morphotype. Some singletons (MFGs with a single morphotype) corresponded to taxa identified at high resolution (genus) and may reflect that those morphologies were informative for taxonomic ID. Conversely, other singletons represented taxa identified at low resolution (phylum, class, order), and may reflect the morphologies were not informative for fine taxonomic ID from images. MFGs comprising more than one morphotype (replicates) were either taxonomically heterogeneous (composed of different taxa sharing similar traits) or taxonomically homogeneous (sharing similar traits and, e.g., all belonging to the order Alcyonacea). Indeed, our results revealed a tight link between the growth forms and the taxonomy particularly for cnidarians (e.g., quill for Pennatuloidea, pinnate for Antipatharia, large fleshy polyps for Actiniaria). Some traits used in the classification were nevertheless inferred from taxonomic knowledge, especially for cnidarians (e.g., root system of *Acanella* or level of skeleton rigidity) and might partly explain this close relationship. However, for cnidarians, the taxonomic IDs have been pushed further than for sponges especially because morphologies were often informative at least for ID to the order rank, although some growth forms can mix different orders (e.g., bottle-brush for Antipatharia and Alcyonacea). Thus, MFGs capture morphological diversity that may reflect differences in structural complexity and potentially associated functions. MFGs also partly reproduce taxonomic diversity, which reflects evolutionary history.

The morpho-functional approach captured a component of the morphological and structural diversity of sponge and cnidarian assemblages that was not represented by the level of taxonomic identification reached on images. Considering the taxonomic dataset was curated for community analysis to keep a balance between richness/abundance and to avoid taxonomic overlap, the morpho-functional approach is also more integrative. It makes it possible to retain a larger proportion of the dataset and thus to be representative of the community’s diversity. The richness difference between the two classifications was particularly notable for sponges, considering we identified only two classes in the taxonomic dataset (Demospongiae, Hexactinellida), but that among these taxa, a high morpho-functional diversity was observable on images. Using image-based taxonomy, one can detect a taxonomic richness constrained by the achievable ID rank, by the sampling effort in the area, and by methodological constraints. Indeed, in many studies, different taxonomic ranks are combined to capture the greatest possible diversity for community analysis (e.g., Girard et al., 2024; Vlach et al., 2025). Even using MFGs, a higher morpho-functional diversity could be expected with a higher sampling effort (as shown by the rarefaction curves in (**Supplem. mat. 7A**).

The morpho-functional classification also revealed a small-scale variability of the sponge and cnidarian morphologies, that was inferred by the number and spatial spacing of the sites with significant LCBD values, i.e., having unique MFG composition. This small-scale variability was not detected using the available taxonomic resolution. These MFGs produced structural complexity at a small spatial scale and created unique biogenic habitats for other potential associated species (Hawkes et al., 2019 ; Meretta et al., 2024). This level of structural complexity was not detected using taxonomy owing to the small number of taxa described. This was especially pronounced for sponges, for which only two taxonomic classes were reached, whereas a much greater number of MFGs contributed to beta-diversity (“SCBD”), and MFGs also captured local differences in unique assemblage composition (“LCBDs”) that were not consistently detected at the taxonomic level. For cnidarians, differences between the two approaches were less pronounced for SCBD, but MFGs nevertheless detected a greater spatial variability in unique assemblages than the taxonomic classification (mostly orders). Those patterns are concordant with the study of Untiedt et al., 2021, showing the loss of information and poor performance of a cnidarian VME classification scheme (i.e., cnidarian orders) for community-level differentiation, compared to morphological one (CATAMI).

Although MFGs cannot be interpreted as direct measures of habitat complexity or functional diversity, their ability to distinguish morphological states that are not resolved taxonomically may therefore provide ecologically relevant information about the distribution of potential biogenic habitats. An in-depth analysis of MFGs’ role in the structure of the surrounding megabenthic community showed that the MFG approach detects significant correlations with benthic assemblages, although the percentage of variance explained is minimal. Conversely, the taxonomic approach showed no significant effect on benthic variance. Although only a small part of the spatial structure of the megafauna community was explained by the occurrence of MFGs after accounting for spatial and environmental variables, the study evidenced some correlations between mobile, or sessile taxa and these biological structures whose size, flexibility, and architectural complexity – some of the characters we used to define the groups – are related to species diversity (Buhl-Mortensen et al., 2010, 2016).

The notion of associated taxa used in this study does not reflect strictly the common sense of associates (i.e., physically living on/inside the host) (Buhl-Mortensen & Buhl-Mortensen, 2004; De Clippele et al., 2015). In the present study, two types of co-structuring were detected. (1) Either the associated fauna was systematically physically present on sponges/cnidarians, or (2) there were co-occurrence with particular morphologies within the same area. For example, the MFG ‘C1’ (including Zoantharia and *Epizoanthus* taxa) was correlated to Hexactinellida and Paguroidea. Image observations confirmed that zoantharian colonies were attached to large stalked sponges (Hyalonematidae) while never observed on hard bottom or sediment, a commonly reported association (Beaulieu, 2001; Kise et al., 2022). It is likely that these zoantharian colonies would benefit from the elevated position of this hard substratum to enhance their filtering activity and to avoid epibenthic predators (Beaulieu, 2001). Similarly, small colonies of *Epizoanthus* zoantharians were repeatedly observed living on paguroids, while the MFG ‘C8’ (small epibenthic actinians) also showed a significant correlation with Paguroidea (an association also observed in the images). The analyses also revealed significant correlations between MFGs and particular taxa in close proximity but without physical associations. For example, we detected multiple correlations between sponges and cnidarians taxa and morphologies, such as between Alcyonaceae and large aerial bushy sponges, and between Zoanthidae and aerial amphoras, small aerial bushy and large aerial stalked cup sponge morphologies, including *Aphrocallistes* sp. Such patterns may reflect shared environmental preferences or responses to environmental conditions that were not fully represented by the variables included in the analyses. They could also result from indirect biotic relationships or from spatially structured processes such as recruitment and dispersal. Finally, we also observed a tendency towards higher densities of some fish taxa (e.g., *Chlorophthalmus*, Lophiiformes, Anguilliformes, Pleuronectiformes), as well as gastropods and brachiopods in areas where some sponge and cnidarian MFGs occurred at higher densities. Previous studies have observed an effect of cold-water corals or sponge habitat on fish abundance or at least a co-structuring (D’Onghia et al., 2010; Linley et al., 2017; Hawkes et al., 2019; Boulard et al., 2023; Coppock et al., 2024). Such relationships may arise because structurally complex habitat-forming organisms provide refuge, attachment surfaces or feeding opportunities, although the mechanisms cannot be inferred from the present data. Smaller benthic organisms have also been reported for sponge and coral habitats, where the three-dimensional structure of the hosts can provide habitat for a diverse associated fauna, including gastropods, amphipods, copepods and polychaetes (Wulff, 2006; Buhl-Mortensen et al., 2010; Pierrejean et al., 2020).

Those are interesting insights into biological interaction/association but should be interpreted cautiously. The very low proportion of explained variance indicates that MFGs account for only a tiny component of the benthic assemblage spatial structure. The large residual fraction may reflect several processes that were not explicitly captured by the analyses, including unmeasured environmental gradients, stochastic variation, recruitment and dispersal processes, and biotic interactions. In addition, many other associations were not visible as we only considered megafauna in the images (the visible biodiversity) while sponge and coral associations with cryptic species, macrofauna, meiofauna and other small-sized organisms commonly occur (e.g., copepods, amphipods, polychaetes) (Wulff, 2006; Buhl-Mortensen et al., 2010; Pierrejean et al., 2020). In this case, image transects are not the most suitable method for studying associations between these organisms which need to be sampled. Nevertheless, these results suggest that the additional structural information captured by MFGs can be relevant for investigating the spatial distribution of potential biogenic habitats and their associated fauna.

The additional morphological resolution provided by MFGs was also reflected in their correlation with environmental gradients especially for sponges. MFG composition for sponges was correlated with a broader set of environmental variables and, in particular, with finer-scale drivers such as geomorphology, topography and substrate, whereas the taxonomic approach was mostly correlated with broad-scale gradients such as depth, current conditions and chlorophyll concentration. The correspondence observed between the environmental associations of Demospongiae and Hexactinellida and those of their respective morphologies suggests that part of the environmental signal is shared between taxonomic identity and morphology. However, the MFG approach further discriminated among morphologies within these broad taxonomic groups. Higher densities of demosponges were observed in environments with strong turbulent currents, while in these hydrodynamic environments, several compact/intermediate forms belonging to the Demospongiae and Porifera (indet.) were correlated to carbonate substrate & geomorphology. Conversely, Hexactinellida were associated with less turbulent water flow, and several hexactinellid morphologies characterised by aerial forms tended to correlate with volcanic, sloping and rougher areas. In our classification, it was mainly Hexactinellida that were associated with the ‘aerial’ trait, although some did not have this trait. The abiotic drivers of the observed sponge morphologies in our studies, regardless of their taxonomy, corroborate the hypothesis reviewed in Schönberg, 2021 (further detailed in the second section of the discussion). For cnidarians in contrast, we did not explain the spatial structures of MFGs in relation to the environment more specifically than with taxonomy. Indeed, we found correspondences between the cnidarian orders and their related morphologies with the environment (set of significant drivers) in which they lived. It seems that the study of the environmental drivers of the cnidarian spatial structures does not require an in-depth classification based on morphology. However, given the paucity of studies looking at the drivers of morphological response of deep-sea cnidarians (De Clippele et al., 2018; Sanna et al., 2023; Sanna, 2024) and the complexity of the taxonomical and morphological overlap for cnidarians, further comparative studies are needed to generalise this conclusion.

### Drivers and ecological interpretation of sponge and cnidarian MFGs spatial structure along the Mozambique Channel

The morpho-functional approach revealed richness differences among sites and spatial variability, occurring even on small spatial scales (e.g., Mayotte island slopes).

The richness of sponge and cnidarian morpho-functional groups differed among the studied seamounts and between seamounts and island slopes in the Mozambique Channel. A previous study on sponge morphological diversity in the Western Indian Ocean revealed a major influence of substratum nature and heterogeneity in explaining sponge morphological diversity (Bell & Barnes, 2001). It is likely that the high level of substrate diversity quantified along Mayotte slopes (Hanafi-Portier et al., 2024) partly explains their higher sponge MFG richness/diversity. A decrease in sponges’ morphological diversity in response to increasing current regime was observed in Barnes & Bell (2002) because of the removal of more fragile forms. In our study, the high to intermediate current speed and turbulence crossing Sakalaves and Hall banks may also select for a small range of sponge morphologies able to support such hydrodynamics (notably compact cups). The Glorieuses platform was composed of a higher diversity of sponge MFGs (e.g., ball, creeping forms). These morphologies have large contact surfaces with the seafloor and are generally favoured by strong currents (Bell & Barnes, 2000; Schönberg, 2021). The high densities forming sponge grounds on the summit of this seamount may be favored by internal waves resulting from the interaction with the topography of the seamounts which allow the resuspension of organic matter and the supply of food (Hanz et al., 2021; Wong et al., 2025). In contrast, the frequent and strong currents could partly explain the low diversity observed in cnidarians MFGs. Such a loss of morphological diversity and structural complexity has been observed with 3D-simulated coral communities (Cresswell et al., 2020), notably explained by the frequency (temporal dynamics) over the intensity of the currents.

Porifera and Cnidaria MFG beta diversity differed also between seamounts and island slopes, and was higher along the island slopes and lower on the flat platforms, both for sponges and cnidarians. This beta diversity was characterised by a spatial distribution of unique assemblages of sponge and cnidarian morphologies, forming biogenic habitats at smaller scales along island slopes compared to seamounts. Several factors can explain these differences, such as the higher substrate diversity, especially over short distances, and the higher - relatively narrow - bathymetric gradient (up to 734 m) along the slopes of Mayotte compared to the flat platforms (∼100 m). Some unique biogenic habitats (assemblages) were thus preferably localised on volcanic areas or at interface between different substrate types and geomorphologies. Similarly, a greater diversity of sponge communities was observed along the slopes than at the summit of Schulz Bank, and a sponge ground zonation appeared to follow a greater depth gradient (∼2000 m) and change with substrate type (Meyer et al., 2023). For sponges, the discrepancy observed at Glorieuses between a relatively high sponge MFGs richness and low beta diversity suggests that, at this site, the various sponge communities vary only marginally with a lot of rare MFGs and dominance of one single MFG. These various sponge MFGs reflect a similar response to the strong but stable current conditions that flow through the platform, which may also contribute to the low beta-diversity.

Sponges heavily rely on local hydrodynamics for survival; coupled with their active pumping to ensure oxygenation and feed on suspended organic matter, strong water flows also facilitate the exchange of gametes and clean their waste, preventing detrimental smothering by high sedimentation (Schönberg, 2021). Amorphous (massive) simple sponges, encountered under the high and stable currents of the Glorieuse platform, are not suited to stagnant water and are at risk of being clogged by sedimentation, so prefer intermediate to high and predictable current velocity (Schönberg, 2021). Although cup-like sponges appear not to be well suited to strong and variable hydrodynamics, they appear abundant under the moderate current conditions - relatively to Glorieuses platform - on the Sakalave platform. Their dominance at this site may be explained by their tolerance to oxygen-reduced environments (Schönberg, 2021), which characterise the overlying waters of the Sakalaves platform (Hanafi-Portier et al., 2024). Compact sponges (e.g., crusts, creeping and amorphous) were also correlated to carbonate rock and geomorphology coverage. It is likely that the high contact surface of these morphologies favours the homogeneous bottom provided by carbonate slab as more robust anchoring, and that the high rocky coverage highlights associated low sedimentary rate, favouring growth forms poorly resistant to sedimentation pressure (e.g., crust) (Schönberg, 2021). Wong et al. (2025) similarly observed a preference of encrusting sponges for relatively low-rugosity seabeds under wave-exposed conditions, suggesting that hydrodynamic and substrate characteristics may jointly influence the distribution of these compact growth forms. At the opposite, delicate sponge morphologies, with aerial compactness (e.g., bushy, stalked and vases) were mostly correlated to topography variables and substrate including volcanic rock or mixed with sediment while were weakly correlated to current variability. These growth forms commonly develop in nutrient-depleted environments, with poor water and nutrient mixing (Schönberg, 2021).

Concerning cnidarians, topography and substrate played a predominant structuring role in shaping their MFGs in the northern Mozambique Channel, compared to hydrological variables (deep current speed and variability and winter Chla concentration) conversely to sponges. However the scale of observation was larger for sponges, with data along the entire channel. With a likely higher contribution of the hydrodynamic processes (i.e., mesoscale eddies) taking place in the centre and south of the channel (Hancke et al., 2014) and therefore driving part of the variability at this scale.

Substrate preferences of cnidarians through their various anchoring modes can explain differences in MFGs composition between and within sites. For example, Pennatuloids have a peduncled anchoring mode, embedded in the sediment (Bayer, 1983), and were correlated to fine sediment observed along the northern and western slope of Mayotte. Conversely, medium to large arborescent, bottle-brush (e.g., Chrysogorgiidae, Alcyonacea) or bushy (e.g., Zoanthidae) cnidarians have a calcified disc as attachment mode, and were correlated to volcanic geomorphology, and to rugosity that characterised roughness provided by volcanic reliefs. These conditions could benefit the settlement of these stalked morphologies. These medium to large stalked colonial morphologies were also correlated to slope. It is known that suspension feeding corals favour sloping areas where rectified flow and current acceleration enhance food particle resuspension and so food fluxes (Genin et al., 1986). We have highlighted a significant role of deep currents, especially their variability, on the structure of cnidarian morpho-functional groups, by differentiating the distribution of some semi-rigid bushy or large arborescent forms under high current variability/speed, as opposed to solitary fleshy and small rigid arborescent forms under low hydrodynamic regimes. Few other studies have revealed a role for deep currents in shaping cold-water coral architectures (De Clippele et al., 2018; Sanna et al., 2023). More compact morphologies would be expected under high current velocity (De Clippele et al., 2018, Sanna et al., 2023), having a bush-like shape with branched growth in the flow direction under unidirectional current (Chindapol et al., 2013; De Clippele et al., 2018). Our results tend to correspond to these expectations, but additional data on the nature of the currents (direction, speed and temporality at a finer resolution) are required, as we also found large non-compact arborescent forms under such a flow regime.

Both for sponge and cnidarian MFGs, the hydrology displayed high shared contribution with topography, substrate and geomorphology in explaining MFGs variance in composition, revealing the complex interaction prevailing between the seamounts topography, rectified current flow and resulting processes in food fluxes and larvae recruitment (Marsac et al., 2020; Dai et al., 2022).

The low variance explained in our analysis is a common observation in ecological studies especially in deep-sea. Some drivers were not measured (nutrients, POM, etc.) and might provide higher explanatory power to the sponge/cnidarian community structure. We also did not consider many other morpho-functional traits not or hardly observable/measurable from images, but having relevance to understand ecosystem functioning (e.g., Cárdenas & Rapp, 2013; Gómez-Gras et al., 2025). It highlights also a spatial mismatch as while our environmental layers (bathymetry, slope) are resolved at a 10-meter grid scale, the biological processes governing sponge and coral larval settlement operate at a centimetric scale of micro-topography. Furthermore, the apparent spatial co-structuring between specific sponge MFGs and mobile associated fauna must be interpreted with caution. While it strongly suggests habitat provisioning, it can also partly reflect a shared ecological response to unmeasured micro-hydrodynamic conditions rather than mandatory biological interactions. We observed a high proportion of variance explained by spatial variables which may include environmental variables spatially structured but not measured, or not identified by analyses (part of the variability is linked to individual taxa forming the MFGs); or biological processes (population dynamics, interactions between taxa, neutral processes) which could be detected for MFGs gathering one or a few taxa. Finally, a better ecological interpretation of the morpho-functional groups delineated from the images and a comparison between the functional *vs.* the taxonomic diversity of the biogenic habitats could be possible by calibrating the images with sampling of physical specimens (e.g., from a ROV) which would allow finer resolution of the taxonomic IDs from images of the individuals and their varying morpho-traits.

### Contribution of the MFG approach and integration into standardised and interoperable frameworks

The proposed method makes it possible to formalise the description of the morphological diversity observed in the images, as independently as possible of the taxonomic ID. This approach offers a compromise that better integrates the structural diversity of habitat forming communities, and their potential functional dimension. This approach is less efficient for corals, as the morpho-functional interpretation hypotheses are still very rudimentary and for which the morphological diversity described was mainly informative of a better taxonomic ID resolution (orders). Conversely, the morpho-functional approach appears more effective for sponges whose taxonomic ID on images is often constrained at low resolution (e.g., class) and for which much more studies have explored the relationships between morphology and the environment (e.g., Cárdenas & Rapp, 2013; Duckworth, 2016; Pineda et al., 2016; Soto et al., 2018; Gökalp et al., 2020; review in Schönberg, 2021).

In our study, MFGs captured additional structural diversity, revealed finer-scale spatial variability in potential biogenic habitats (higher beta diversity), and identified associations between particular morphologies and the surrounding benthic assemblage. This additional information was particularly relevant for sponges, for which taxonomic identification from imagery was often limited to broad taxonomic levels, whereas substantial morphological variation occurred within these groups. For cnidarians, the gain was more limited because morphological and taxonomic variation were more closely related and because some morphological traits were themselves informed by taxonomic knowledge. The usefulness of the MFG approach therefore depends on both the taxonomic resolution achievable from imagery and the extent to which morphological variation provides additional information beyond taxonomy.

Considering the need for scientific community to converge toward standardisation of image-based data (Bowden et al., 2020; Borremans et al., 2024), the MFG approach in the present study could be translated and integrated into existing hierarchical image-scoring systems such as CATAMI (Althaus et al., 2015). The hierarchical structure of such systems allows observations to be retained at a fine resolution while remaining aggregable to broader, comparable categories across studies. Additional descriptors could be incorporated as nested levels within this structure, particularly for sponges. Traits such as size, compactness or additional visible structural features (e.g., hispidity or outgrowths) could provide information on the structural heterogeneity of habitat-forming taxa and its variation along environmental gradients. Such an approach is consistent with Perkins et al. (2025), who incorporated size descriptors into a CATAMI-based classification and showed that size information provided complementary information to abundance for characterising ecosystem dynamics. Colour may be less suitable for standardised scoring because its apparent expression can vary with image illumination and quality. More generally, descriptors included in such a framework should be observable or measurable consistently across images. For instance, polyp size may vary depending on the degree of polyp retraction at the time of image acquisition, making this trait difficult to measure consistently. Although polyp size helped discriminate Actiniaria from other corals in our study, we would therefore not recommend its use as a standardised descriptor without further assessment of its reproducibility across imaging conditions.

Finally, although sponges and cnidarians were treated separately here, because the relevant morphological traits differed and because we aimed to compare the contribution of the approach between groups, both represent important habitat-forming taxa. Developing a set of common descriptors across habitat-forming groups, complemented by group-specific traits where necessary, could therefore further improve the interoperability of image-based studies. This perspective is consistent with the framework of Benoit et al. (2026), which proposes standardised trait categories for marine foundation species. Embedding such common descriptors within hierarchical systems such as CATAMI could provide a way to retain fine-scale structural information while maintaining comparability across taxa, studies and spatial scales.

## Conclusion

When fine resolution taxonomic ID of sponges and cnidarians is not achievable from imagery, a common encountered problem especially in poorly sampled areas, a morpho-functional approach provides a practical way to retain additional ecological information from image-based surveys. By describing morphological variation through observable traits, MFGs captured structural diversity and spatial patterns of potential biogenic habitats that were not consistently represented by the available taxonomic resolution. They also revealed associations between particular morphologies and other megabenthic taxa, although these relationships explained only a tiny proportion of community variation, and should be interpreted as partial components of broader ecological patterns.

The approach was particularly informative for sponges, for which morphology provided information beyond the coarse taxonomic resolution achievable from imagery and clear relationships with the environment. Sponge morphologies respond well to the environment, and, with a good understanding of these relationships, offer the possibility to predict their distribution and that of the habitat they form using environmental proxies. Inversely, other studies have proposed using sponge morphologies as proxies for the environmental conditions in which they are found (i.e., ecological indicators) (Mary George et al., 2018; Schönberg, 2021). However, the low explanatory power of the environmental variables considered here also indicates that additional, unmeasured environmental, ecological or historical processes contribute to the distribution of sponge morphologies, as well as potential scale mismatches between the spatial resolution of available environmental variables and the fine-scale patterns of faunal structure.

For cnidarians however, the additional information provided by MFGs was more limited, and a combined approach using taxonomic entities (to the achievable rank) and gross morphology could be a reasonable trade-off (Untiedt et al., 2021).

An important feature of our approach was the use of multiple morphological traits combined with numerical classification to reduce subjectivity in the delineation of morphological groups and to identify replicate within morphological combinations. This framework could nevertheless be simplified and standardised by integrating such information directly into existing hierarchical image-scoring systems such as CATAMI (Althaus et al., 2015). This way, instead of study-specific MFG schemes, predefined morphological categories could be complemented by nested trait descriptors providing finer structural resolution. For sponges, these could include size, compactness, and visible external features such as hispidity or outgrowths. Such a hierarchical framework would preserve comparability at broader levels while allowing more detailed morphological categories to be used as variables in ecological analyses.

To our knowledge, this is the first study applying beta diversity metrics, including quantitative assessment of ‘taxa’ (SCBD) or sites (LCBD) contributions to beta diversity, to morpho-functional groups. Applying these metrics to MFGs provide a quantitative framework to identify vulnerable habitats (from SCBD metrics) and their spatial scales of variability (from LCBD metrics), performing different potential functional roles and responding differently to disturbance according to their morpho-functional groups/traits composition. Complementary to taxonomic information, the morphological traits observable and assessed from imagery provide useful information regarding the ecological function, life history and vulnerability of sponges and cnidarians. Frameworks based on morphologies could therefore also be considered in the recent proposition of VME designation from imagery (Baco et al., 2023).

## Acknowledgements

We acknowledge the chief scientists and scientific team of the BIOMAGLO (Laure Corbari, Sarah Samadi, Karine Olu, 2017, https://doi.org/10.17600/17004000), BATHYMAY (Pol Guennoc, 2004, https://doi.org/10.17600/4200020), PTOLEMEE (Stefan Jorry, 2014, https://doi.org/10.17600/14000900), PAMELA-MOZ01 (Karine Olu, 2014, https://doi.org/10.17600/14001000), and PAMELA MOZ04 (Gwénaël Jouet and Éric Deville, 2015, https://doi.org/10.17600/15000700) cruises as well as the PAMELA project leaders (Jean François Bourillet, Philippe Bourges, and Jean-Noël Ferry, https://doi.org/10.18142/236). The authors are grateful to the captains and crew of RV’s *L’Antea*, *L’Atalante*, *Le Pourquoi Pas?* as well as to the SCAMPI towed-camera team for their contribution to field data acquisition. We also thank all the engineers who contributed to the project: Marie Eve-Julie Pernet for handling the faunal physical samples used as voucher specimens for image-based faunal identification and Louise Keszler for the primary annotation of the BIOMAGLO campaign images, Julie Tourolle for assistance with GIS data, Catherine Borremans and Olivier Soubigou for their assistance with image annotations and GIS development needs, Christophe Brandily for assistance with CTD data processing, and Charline Guerin for reprocessing the bathymetric resolution of the DTM. We also thank Simon Gourdon for assistance in the morphotype and substrate annotations from images, and Simon Courgeon for assistance with the interpretation of geomorphological units. We are grateful to Tim W. Nattkemper and Daniel Langenkämper for enabling image storage and their technical assistance using Biigle 2.0. Finally, we thank all the taxonomists involved in the project for their help in identifying fauna from the images: Nicolas Puillandre (gastropods); Philippe Maestrati (bivalves); Éric Pante and Daniela Pica (cnidarians); Paco Cárdenas and Cécile Debitus (sponges); Enrique Macpherson (galatheids); Tin-Yam Chan (crustaceans); Wei-Jen Chen, Jhen-Nien Chen, Mao-Ying Lee and Paul Giannasi (fishes); Thomas Saucède (echinoids) and Christopher Mah (asteroids).

## Data, scripts, code, and supplementary information availability

Data, code and supplementary information are available online: https://doi.org/10.17882/108256 (Hanafi-Portier et al., 2025).

## Conflict of interest disclosure

The authors declare that they comply with the PCI rule of having no financial conflicts of interest in relation to the content of the article. The authors declare the following non-financial conflict of interest: Eric Pante is a recommender for PCI Evolutionary Biology and PCI Genomics.

## Funding

The PAMELA project (PAssive Margin Exploration LAboratories) is a scientific project led by IFREMER and TotalEnergies in partnership with Université de Bretagne Occidentale, Université Rennes 1, Université Pierre et Marie Curie, CNRS and IFPEN. The cruise and the BIOMAGLO project received funding from the Xth European Development Fund (Fonds Européen de Développement ; FED) ‘sustainable management of the natural heritage of Mayotte and the Eparses Islands’ program led by the French Southern and Antarctic Lands (Terres Australes et Antarctiques Françaises ; TAAF) with the support of the Mayotte Departmental Council (Conseil Départemental de Mayotte), the French Development Agency (Agence Française de Développement ; AFD) and the European Union. The thesis of Mélissa Hanafi-Portier is co-funded by TotalEnergies and IFREMER as part of the PAMELA (Passive Margin Exploration Laboratories) scientific project.

## Notes

https://doi.org/10.17882/108256

### Summary of Updates

The main changes made in this revised version include: - A complete revision of the Results section, including a direct comparison between the MFG-based and taxonomic approaches for both sponges and cnidarians. - A revision of the Discussion, including a new section highlighting the contribution and limitations of the MFG approach. In particular, we further discuss the potential of using morphology as a reproducible framework for comparing assemblages across studies, and the possibility of transferring this framework to a standardized classification system such as CATAMI, including suggestions for potential additions to the CATAMI classification. We also provide a more detailed discussion of the relatively low proportion of variance explained by the models. - A revision of the figures, including modifications to Figure 1 (maps), removal of the former Figure 2, and the addition of new figures presenting pairwise comparisons between the MFG-based and taxonomic approaches. - Additional analyses, including Venn diagrams to assess the potential structuring effect of sponges and cnidarians on the surrounding megabenthic community.

## References

1. Althaus, F., Hill, N., Ferrari, R., Edwards, L., Przeslawski, R., Schönberg, C. H. L., Stuart-Smith, R., Barrett, N., Edgar, G., Colquhoun, J., Tran, M., Jordan, A., Rees, T., & Gowlett-Holmes, K. (2015). A Standardised Vocabulary for Identifying Benthic Biota and Substrata from Underwater Imagery : The CATAMI Classification Scheme. PLOS ONE, 10(10), e0141039. 10.1371/journal.pone.0141039

2. Audru, J.-C., Guennoc, P., Thinon, I., & Abellard, O. (2006). Bathymay : La structure sous-marine de Mayotte révélée par l’imagerie multifaisceaux. Comptes Rendus Geoscience, 338(16), 1240–1249. 10.1016/j.crte.2006.07.010

3. Baco, A. R., Morgan, N. B., & Roark, E. B. (2020). Observations of vulnerable marine ecosystems and significant adverse impacts on high seas seamounts of the northwestern Hawaiian Ridge and Emperor Seamount Chain. Marine Policy, 115, 103834. 10.1016/j.marpol.2020.103834

4. Baco, A. R., Ross, R., Althaus, F., Amon, D., Bridges, A. E. H., Brix, S., Buhl-Mortensen, P., Colaco, A., Carreiro-Silva, M., Clark, M. R., Du Preez, C., Franken, M.-L., Gianni, M., Gonzalez-Mirelis, G., Hourigan, T., Howell, K., Levin, L. A., Lindsay, D. J., Molodtsova, T. N., … Yesson, C. (2023). Towards a scientific community consensus on designating Vulnerable Marine Ecosystems from imagery. PeerJ, 11, e16024. 10.7717/peerj.16024

5. Barnes, D. K. A., & Bell, J. J. (2002). Coastal sponge communities of the West Indian Ocean : Morphological richness and diversity. African Journal of Ecology, 40(4), 350–359. 10.1046/j.1365-2028.2002.00388.x

6. Bayer, F. M. (Éd.). (1983). Illustrated trilingual glossary of morphological and anatomical terms applied to octocorallia. Brill.

7. Beaulieu, S. E. (2001). Life on glass houses : Sponge stalk communities in the deep sea. Marine Biology, 138(4), 803–817. 10.1007/s002270000500

8. Beazley, L. I., Kenchington, E. L., Murillo, F. J., & Sacau, M. del M. (2013). Deep-sea sponge grounds enhance diversity and abundance of epibenthic megafauna in the Northwest Atlantic. ICES Journal of Marine Science, 70(7), 1471–1490. 10.1093/icesjms/fst124

9. Bell, J. (2007). Contrasting patterns of species and functional composition of coral reef sponge assemblages. Marine Ecology Progress Series, 339, 73–81. 10.3354/meps339073

10. Bell, J. B., Alt, C. H. S., & Jones, D. O. B. (2016). Benthic megafauna on steep slopes at the Northern Mid-Atlantic Ridge. Marine Ecology, 37(6), 1290–1302. 10.1111/maec.12319

11. Bell, J. J. (2008). The functional roles of marine sponges. Estuarine, Coastal and Shelf Science, 79(3), 341–353. 10.1016/j.ecss.2008.05.002

12. Bell, J. J., & Barnes, D. K. A. (2000). The influences of bathymetry and flow regime upon the morphology of sublittoral sponge communities. Journal of the Marine Biological Association of the United Kingdom, 80(4), 707–718. 10.1017/S0025315400002538

13. Bell, J. J., & Barnes, D. K. A. (2001). Sponge morphological diversity : A qualitative predictor of species diversity? Aquatic Conservation: Marine and Freshwater Ecosystems, 11(2), 109–121. 10.1002/aqc.436

14. Bell, J. J., & Barnes, D. K. A. (2002). Modelling sponge species diversity using a morphological predictor : A tropical test of a temperate model. Journal for Nature Conservation, 10(1), 41–50. 10.1078/1617-1381-00005

15. Bo, M., Bertolino, M., Bavestrello, G., Canese, S., Giusti, M., Angiolillo, M., Pansini, M., & Taviani, M. (2012). Role of deep sponge grounds in the Mediterranean Sea : A case study in southern Italy. Hydrobiologia, 687(1), 163–177. 10.1007/s10750-011-0964-1

16. Borcard, D., Gillet, F., & Legendre, P. (2018). Numerical Ecology with R. Springer International Publishing. 10.1007/978-3-319-71404-2

17. Borremans, C., Durden, J., Schoening, T., Curtis, E., Adams, L., Branzan Albu, A., Arnaubec, A., Ayata, S.-D., Baburaj, R., Bassin, C., Beck, M., Bigham, K., Boschen-Rose, R., Collett, C., Contini, M., Correa, P., Dominguez-Carrió, C., Dreyfus, G., Duncan, G., … Zurowietz, M. (2024). Report on the Marine Imaging Workshop 2022. Research Ideas and Outcomes, 10, e119782. 10.3897/rio.10.e119782

18. Boulard, M., Lawton, P., Baker, K., & Edinger, E. (2023). The effect of small-scale habitat features on groundfish density in deep-sea soft-bottom ecosystems. Deep Sea Research Part I: Oceanographic Research Papers, 193, 103891. 10.1016/j.dsr.2022.103891

19. Bourillet, J.-F., Ferry, J.-N., & Bourges, P. (2013). PAMELA : PASSIVE MARGINS EXPLORATION LABORATORIES. 10.18142/236

20. Bowden, D. A., Rowden, A. A., Chin, C. C., Hempel, S., Wood, B. A., Hart, A. C., Clark, M. R., & Fisheries New Zealand (Government agency). (2020). Best practice in seabed image analysis for determining taxa, habitat, or substrata distributions. https://fs.fish.govt.nz/Doc/24797/AEBR-239-Best-Practice-In-Seabed-Image-Analysis.pdf.ashx

21. Buhl-Mortensen, L., & Buhl-Mortensen, P. (2004). The distribution and diversity of species associated with deep sea gorgonian corals off the Atlantic Canada. 849–879. 10.1007/3-540-27673-4_44

22. Buhl-Mortensen, L., Vanreusel, A., Gooday, A. J., Levin, L. A., Priede, I. G., Buhl-Mortensen, P., Gheerardyn, H., King, N. J., & Raes, M. (2010). Biological structures as a source of habitat heterogeneity and biodiversity on the deep ocean margins : Biological structures and biodiversity. Marine Ecology, 31(1), 21–50. 10.1111/j.1439-0485.2010.00359.x

23. Buhl-Mortensen, P., Buhl-Mortensen, L., & Purser, A. (2016). Trophic Ecology and Habitat Provision in Cold-Water Coral Ecosystems. In S. Rossi, L. Bramanti, A. Gori, & C. Orejas (Éds.), Marine Animal Forests (p. 1–26). Springer International Publishing. 10.1007/978-3-319-17001-5_20-1

24. Cairns, S. D. (2007). Deep-water corals : An overview with special reference to diversity and distribution. BULLETIN OF MARINE SCIENCE, 81(3).

25. Cárdenas, P., & Rapp, H. (2013). Disrupted spiculogenesis in deep-water Geodiidae (Porifera, Demospongiae) growing in shallow waters. Invertebrate Biology, 132. 10.1111/ivb.12027

26. Chappell, J. (1980). Coral morphology, diversity and reef growth. Nature, 286(5770), 249–252. 10.1038/286249a0

27. Chindapol, N., Kaandorp, J. A., Cronemberger, C., Mass, T., & Genin, A. (2013). Modelling Growth and Form of the Scleractinian Coral Pocillopora verrucosa and the Influence of Hydrodynamics. PLoS Computational Biology, 9(1), e1002849. 10.1371/journal.pcbi.1002849

28. Coppock, A. G., Kingsford, M. J., & Jones, G. P. (2024). Importance of complex sponges as habitat and feeding substrata for coral reef fishes. Marine Biology, 171(8), 154. 10.1007/s00227-024-04467-6

29. Corbari, L., Samadi, S., & Olu, K. (2017). BIOMAGLO cruise, RV Antea. 10.17600/17004000

30. Courgeon, S., Jorry, S. J., Camoin, G. F., BouDagher-Fadel, M. K., Jouet, G., Révillon, S., Bachèlery, P., Pelleter, E., Borgomano, J., Poli, E., & Droxler, A. W. (2016). Growth and demise of Cenozoic isolated carbonate platforms : New insights from the Mozambique Channel seamounts (SW Indian Ocean). Marine Geology, 380, 90–105. 10.1016/j.margeo.2016.07.006

31. Cresswell, A. K., Thomson, D. P., Haywood, M. D. E., & Renton, M. (2020). Frequent hydrodynamic disturbances decrease the morphological diversity and structural complexity of 3D simulated coral communities. Coral Reefs, 39(4), 1147–1161. 10.1007/s00338-020-01947-1

32. Dai, S., Zhao, Y., Li, X., Wang, Z., Zhu, M., Liang, J., Liu, H., & Sun, X.-X. (2022). Seamount effect on phytoplankton biomass and community above a deep seamount in the tropical western Pacific. Marine Pollution Bulletin, 175, 113354. 10.1016/j.marpolbul.2022.113354

33. De Clippele, L. H., Buhl-Mortensen, P., & Buhl-Mortensen, L. (2015). Fauna associated with cold water gorgonians and sea pens. Continental Shelf Research, 105, 67–78. 10.1016/j.csr.2015.06.007

34. De Clippele, L. H., Huvenne, V. A. I., Orejas, C., Lundälv, T., Fox, A., Hennige, S. J., & Roberts, J. M. (2018). The effect of local hydrodynamics on the spatial extent and morphology of cold-water coral habitats at Tisler Reef, Norway. Coral Reefs, 37(1), 253–266. 10.1007/s00338-017-1653-y

35. Denis, V., Ribas-Deulofeu, L., Sturaro, N., Kuo, C.-Y., & Chen, C. A. (2017). A functional approach to the structural complexity of coral assemblages based on colony morphological features. Scientific Reports, 7(1). 10.1038/s41598-017-10334-w

36. D’Onghia, G., Maiorano, P., Sion, L., Giove, A., Capezzuto, F., Carlucci, R., & Tursi, A. (2010). Effects of deep-water coral banks on the abundance and size structure of the megafauna in the Mediterranean Sea. *Deep Sea Research Part II: Topical Studies in Oceanography, The APLABES Programme: Physical*, Chemical and Biological Characterization of Deep-Water Coral Ecosystems from the Ionian Sea (Mediterranean), 57(5), 397–411. 10.1016/j.dsr2.2009.08.022

37. Dray, S., Bauman, D., Blanchet, G., Borcard, D., Clappe, S., Guenard, G., Jombart, T., Larocque, G., Legendre, P., Madi, N., & Wagner, H. H. (2022). *Adespatial : Multivariate Multiscale Spatial Analysis. R package version 0.3-16* [Logiciel]. https://CRAN.R-project.org/package=adespatial

38. Dray, S., & Dufour, A.-B. (2007). The ade4 Package : Implementing the Duality Diagram for Ecologists. Journal of Statistical Software, 22(4). 10.18637/jss.v022.i04

38a. Duckworth, A. R. (2016). Substrate type affects the abundance and size of a coral-reef sponge between depths. Marine and Freshwater Research, 67(2), 246. 10.1071/MF14308

39. FAO (Éd.). (2009). International guidelines for the management of deep-sea fisheries in the high seas : = Directives internationales sur la gestion de la pêche profonde en haute mer. Food and Agriculture Organization of the United Nations.

40. Genin, A., Dayton, P. K., Lonsdale, P. F., & Spiess, F. N. (1986). Corals on seamount peaks provide evidence of current acceleration over deep-sea topography. Nature, 322(6074), 59–61. 10.1038/322059a0

41. Girard, F., Caress, D., Paduan, J., Kuhnz, L., Litvin, S., Flattery, E., Kahn, A., DeVogelaere, A., Burton, E., Lovera, C., Martin, E., & Barry, J. (2024). Habitat heterogeneity over multiple scales supports dense and diverse megafaunal communities on a northeast Pacific ridge. Limnology and Oceanography, 70, 377–392. 10.1002/lno.12766

42. Gökalp, M., Kooistra, T., Rocha, M. S., Silva, T. H., Osinga, R., Murk, A. J., & Wijgerde, T. (2020). The Effect of Depth on the Morphology, Bacterial Clearance, and Respiration of the Mediterranean Sponge Chondrosia reniformis (Nardo, 1847). Marine Drugs, 18(7), Article 7. 10.3390/md18070358

43. Gómez-Gras, D., Linares, C., Viladrich, N., Zentner, Y., Grinyó, J., Gori, A., McFadden, C. S., Fabricius, K. E., & Madin, J. S. (2025). The Octocoral Trait Database : A global database of trait information for octocoral species. Scientific Data, 12(1), 82. 10.1038/s41597-024-04307-8

44. Hadi, T. A., Budiyanto, A., & Wentao, N. (2015). The Morphological and Species Diversity of Sponges in Coral Reef Ecosystem in the Lembeh Strait, Bitung. 13.

45. Hanafi-Portier, M., Samadi, S., Corbari, L., Boulard, M., Miramontes, E., Penven, P., Leroy, B., Napoléon, T., Jorry, S. J., & Olu, K. (2024). Multiscale spatial patterns and environmental drivers of seamount and island slope megafaunal assemblages along the Mozambique channel. Deep Sea Research Part I: Oceanographic Research Papers, 203, 104198. 10.1016/j.dsr.2023.104198

46. Hanafi-Portier, M., Samadi, S., Corbari, L., Chan, T.-Y., Chen, W.-J., Chen, J.-N., Lee, M.-Y., Mah, C., Saucède, T., Borremans, C., & Olu, K. (2021). When Imagery and Physical Sampling Work Together : Toward an Integrative Methodology of Deep-Sea Image-Based Megafauna Identification. Frontiers in Marine Science, 8, 749078. 10.3389/fmars.2021.749078

47. Hancke, L., Roberts, M. J., & Ternon, J. F. (2014). Surface drifter trajectories highlight flow pathways in the Mozambique Channel. Deep Sea Research Part II: Topical Studies in Oceanography, 100, 27–37. 10.1016/j.dsr2.2013.10.014

48. Hanz, U., Roberts, E. M., Duineveld, G., Davies, A., van Haren, H., Rapp, H. T., Reichart, G., & Mienis, F. (2021). Long-term Observations Reveal Environmental Conditions and Food Supply Mechanisms at an Arctic Deep-Sea Sponge Ground. Journal of Geophysical Research: Oceans, 126(3). 10.1029/2020JC016776

49. Hawkes, N., Korabik, M., Beazley, L., Rapp, H., Xavier, J., & Kenchington, E. (2019). Glass sponge grounds on the Scotian Shelf and their associated biodiversity. Marine Ecology Progress Series, 614, 91–109. 10.3354/meps12903

50. Hein, J. R., Conrad, T. A., & Staudigel, H. (2010). Seamount mineral deposits, a source of rare metals for high-technology industries. Oceanography, 23(1), 184–189.

51. Hooper, J., & van Soest, R. (2002). Systema Porifera, a guide to the classification of the sponges (in 2 volumes).

52. Howell, K. L., Davies, J. S., Allcock, A. L., Braga-Henriques, A., Buhl-Mortensen, P., Carreiro-Silva, M., Dominguez-Carrió, C., Durden, J. M., Foster, N. L., Game, C. A., Hitchin, B., Horton, T., Hosking, B., Jones, D. O. B., Mah, C., Laguionie Marchais, C., Menot, L., Morato, T., Pearman, T. R. R., … Wagner, D. (2019). A framework for the development of a global standardised marine taxon reference image database (SMarTaR-ID) to support image-based analyses. PLOS ONE, 14(12), e0218904. 10.1371/journal.pone.0218904

53. James, L., Marzloff, M., Barrett, N., Friedman, A., & Johnson, C. (2017). Changes in deep reef benthic community composition across a latitudinal and environmental gradient in temperate Eastern Australia. Marine Ecology Progress Series, 565, 35–52. 10.3354/meps11989

54. Jouet, G., & Deville, E. (2015). PAMELA-MOZ04 cruise, RV Pourquoi pas ? 10.17600/15000700

55. Kise, H., Montenegro, J., Santos, M. E. A., Hoeksema, B. W., Ekins, M., Ise, Y., Higashiji, T., Fernandez-Silva, I., & Reimer, J. D. (2022). Evolution and phylogeny of glass-sponge-associated zoantharians, with a description of two new genera and three new species. Zoological Journal of the Linnean Society, 194(1), 323–347. 10.1093/zoolinnean/zlab068

56. Kornder, N. A., Esser, Y., Stoupin, D., Leys, S. P., Mueller, B., Vermeij, M. J. A., Huisman, J., & de Goeij, J. M. (2022). Sponges sneeze mucus to shed particulate waste from their seawater inlet pores. Current Biology, 32(17), 3855–3861.e3. 10.1016/j.cub.2022.07.017

57. Kramer, N., Tamir, R., Eyal, G., & Loya, Y. (2020). Coral Morphology Portrays the Spatial Distribution and Population Size-Structure Along a 5–100 m Depth Gradient. Frontiers in Marine Science, 7. 10.3389/fmars.2020.00615

58. Krell, F.-T. (2004). Parataxonomy vs. Taxonomy in biodiversity studies – pitfalls and applicability of ‘morphospecies’ sorting. Biodiversity and Conservation, 13(4), 795–812. 10.1023/B:BIOC.0000011727.53780.63

59. Kruk, C., Huszar, V. L. M., Peeters, E. T. H. M., Bonilla, S., Costa, L., Lürling, M., Reynolds, C. S., & Scheffer, M. (2010). A morphological classification capturing functional variation in phytoplankton. Freshwater Biology, 55(3), 614–627. 10.1111/j.1365-2427.2009.02298.x

60. Kruk, C., Peeters, E. T. H. M., Van Nes, E. H., Huszar, V. L. M., Costa, L. S., & Scheffer, M. (2011). Phytoplankton community composition can be predicted best in terms of morphological groups. Limnology and Oceanography, 56(1), 110–118. 10.4319/lo.2011.56.1.0110

61. Langenkämper, D., Zurowietz, M., Schoening, T., & Nattkemper, T. W. (2017). BIIGLE 2.0—Browsing and Annotating Large Marine Image Collections. Frontiers in Marine Science, 4. 10.3389/fmars.2017.00083

62. Legendre, P., & De Cáceres, M. (2013). Beta diversity as the variance of community data : Dissimilarity coefficients and partitioning. Ecology Letters, 16(8), 951–963. 10.1111/ele.12141

63. Linley, T. D., Lavaleye, M., Maiorano, P., Bergman, M., Capezzuto, F., Cousins, N. J., D’Onghia, G., Duineveld, G., Shields, M. A., Sion, L., Tursi, A., & Priede, I. G. (2017). Effects of cold-water corals on fish diversity and density (European continental margin : Arctic, NE Atlantic and Mediterranean Sea): Data from three baited lander systems. Deep Sea Research Part II: Topical Studies in Oceanography, Towards ecosystem based management and monitoring of the deep Mediterranean, North-East Atlantic and Beyond, 145, 8–21. 10.1016/j.dsr2.2015.12.003

64. Marsac, F., Annasawmy, P., Noyon, M., Demarcq, H., Soria, M., Rabearisoa, N., Bach, P., Cherel, Y., Grelet, J., & Romanov, E. (2020). Seamount effect on circulation and distribution of ocean taxa in the vicinity of La Pérouse, a shallow seamount in the southwestern Indian Ocean. Deep Sea Research Part II: Topical Studies in Oceanography, 176, 104806. 10.1016/j.dsr2.2020.104806

65. Mary George, A., Brodie, J., Daniell, J., Capper, A., & Jonker, M. (2018). Can sponge morphologies act as environmental proxies to biophysical factors in the Great Barrier Reef, Australia? Ecological Indicators, 93, 1152–1162. 10.1016/j.ecolind.2018.06.016

66. McDonald, J. I., Hooper, J. N. A., & McGuinness, K. A. (2002). Environmentally influenced variability in the morphology of Cinachyrella australiensis (Carter 1886) (Porifera : Spirophorida : Tetillidae). Marine and Freshwater Research, 53(1), 79–84. 10.1071/mf00153

67. McFadden, C. S., Ofwegen, L. P. van, & Quattrini, A. M. (2022). Revisionary systematics of Octocorallia (Cnidaria : Anthozoa) guided by phylogenomics. Bulletin of the Society of Systematic Biologists, 1(3), Article 3. 10.18061/bssb.v1i3.8735

68. Meretta, P. E., Sirito, S. M., Schejter, L., Tapia, A. P., Bremec, C., & Genzano, G. (2024). Sea fans and their associated fauna in the marine animal forests of Burdwood Bank at 54°S, SW Atlantic Ocean. Marine Biodiversity, 54(6), 79. 10.1007/s12526-024-01471-3

69. Meroz-Fine, E., Shefer, S., & Ilan, M. (2005). Changes in morphology and physiology of an East Mediterranean sponge in different habitats. Marine Biology, 147(1), 243–250. 10.1007/s00227-004-1532-2

70. Meyer, H., Davies, A., Roberts, E., Xavier, J., Ribeiro, P. A., Glenner, H., Birkely, S.-R., & Rapp, H. (2023). Beyond the tip of the seamount : Distinct megabenthic communities found beyond the charismatic summit sponge ground on an arctic seamount (Schulz Bank, Arctic Mid-Ocean Ridge). Deep Sea Research Part I: Oceanographic Research Papers, 191, 103920. 10.1016/j.dsr.2022.103920

71. Morgan, N. B., & Baco, A. R. (2021). Recent fishing footprint of the high-seas bottom trawl fisheries on the Northwestern Hawaiian Ridge and Emperor Seamount Chain : A finer-scale approach to a large-scale issue. Ecological Indicators, 121, 107051. 10.1016/j.ecolind.2020.107051

72. Oksanen, J., Blanchet, F. G., Friendly, M., Kindt, R., Legendre, P., McGlinn, D., Minchin, P. R., O’Hara, R. B., Gavin, L. S., Peter, S., H. Stevens, M. henry, Szoecs, E., & Wagner, H. (2020). *Vegan : Community Ecology Package. R package version 2.5-7* [Logiciel]. https://CRAN.R-project.org/package=vega

73. Olu, K. (2014). PAMELA-MOZ01 cruise, RV L’Atalante. 10.17600/14001000

74. Pante, E., France, S. C., Couloux, A., Cruaud, C., McFadden, C. S., Samadi, S., & Watling, L. (2012). Deep-Sea Origin and In-Situ Diversification of Chrysogorgiid Octocorals. PLoS ONE, 7(6), e38357. 10.1371/journal.pone.0038357

75. Patton, W. K. (1972). STUDIES ON THE ANIMAL SYMBIONTS OF THE GORGONIAN CORAL, LEPTOGORGIA VIRGULATA (LAMARCK). Bulletin of Marine Science, 22, 419–431.

76. Paz-García, D. A., Aldana-Moreno, A., Cabral-Tena, R. A., García-De-León, F. J., Hellberg, M. E., & Balart, E. F. (2015). Morphological variation and different branch modularity across contrasting flow conditions in dominant Pocillopora reef-building corals. Oecologia, 178(1), 207–218. 10.1007/s00442-014-3199-9

77. Perkins, N. R., Monk, J., Wong, R. H. X., & Barrett, N. S. (2025). Temporal variability in temperate mesophotic ecosystems revealed with over a decade of monitoring with an autonomous underwater vehicle. Marine Environmental Research, 209, 107179. 10.1016/j.marenvres.2025.107179

78. Pica, D., Calcinai, B., Poliseno, A., Trainito, E., & Cerrano, C. (2018). Distribution and phenotypic variability of the Mediterranean gorgonian Paramuricea macrospina (Cnidaria : Octocorallia). The European Zoological Journal, 85(1), 392–408. 10.1080/24750263.2018.1529202

79. Pierrejean, M., Grant, C., Neves, B. de M., Chaillou, G., Edinger, E., Blanchet, F. G., Maps, F., Nozais, C., & Archambault, P. (2020). Influence of Deep-Water Corals and Sponge Gardens on Infaunal Community Composition and Ecosystem Functioning in the Eastern Canadian Arctic. Frontiers in Marine Science, 7, 495. 10.3389/fmars.2020.00495

80. Pineda, M. C., Duckworth, A., & Webster, N. (2016). Appearance matters : Sedimentation effects on different sponge morphologies. Journal of the Marine Biological Association of the United Kingdom, 96(2), 481–492. 10.1017/S0025315414001787

81. Pronzato, R., Bavestrello, G., & Cerrano, C. (1998). MORPHO-FUNCTIONAL ADAPTATIONS OF THREE SPECIES OF SPONGIA (PORIFERA, DEMOSPONGIAE) FROM A MEDITERRANEAN VERTICAL CLIFF. BULLETIN OF MARINE SCIENCE, 63(2), 12.

82. Quattrini, A. M., Ross, S. W., Carlson, M. C. T., & Nizinski, M. S. (2012). Megafaunal-habitat associations at a deep-sea coral mound off North Carolina, USA. Marine Biology, 159(5), 1079–1094. 10.1007/s00227-012-1888-7

83. R Core Team. (2021). R: A language and environment for statistical computing. *Vienna* : *R Foundation for Statistical Computing*.

84. Rogers, A., Blanchard, J. L., & Mumby, P. J. (2014). Vulnerability of Coral Reef Fisheries to a Loss of Structural Complexity. Current Biology, 24(9), 1000–1005. 10.1016/j.cub.2014.03.026

85. Rossi, S., & Bramanti, L. (Éds.). (2020). Perspectives on the Marine Animal Forests of the World. Springer International Publishing. 10.1007/978-3-030-57054-5

86. Rossi, S., Bramanti, L., Gori, A., & Orejas, C. (Éds.). (2017). Marine Animal Forests : The Ecology of Benthic Biodiversity Hotspots. Springer International Publishing. 10.1007/978-3-319-21012-4

87. Sanna, G. (2024). Cold-water coral sensitivity : Growth, morphology and reef-forming potential of Desmophyllum pertusum in response to environmental factors. 10.26092/elib/3389

88. Sanna, G., Büscher, J. V., & Freiwald, A. (2023). Cold-water coral framework architecture is selectively shaped by bottom current flow. Coral Reefs, 42(2), 483–495. 10.1007/s00338-023-02361-z

89. Schönberg, C. H. L. (2016). Happy relationships between marine sponges and sediments – a review and some observations from Australia. Journal of the Marine Biological Association of the United Kingdom, 96(2), 493–514. 10.1017/S0025315415001411

90. Schönberg, C. H. L. (2021). No taxonomy needed : Sponge functional morphologies inform about environmental conditions. Ecological Indicators, 129. 10.101BDj.ecolind.2021.107A0B

91. Shaffer, M. R., Davy, S. K., & Bell, J. J. (2019). Hidden diversity in the genus Tethya : Comparing molecular and morphological techniques for species identification. Heredity, 122(3), 354–369. 10.1038/s41437-018-0134-6

92. Soto, D., Palmas, S. D., Ho, M. J., Denis, V., & Chen, C. A. (2018). Spatial variation in the morphological traits of Pocillopora verrucosa along a depth gradient in Taiwan. PLOS ONE, 13(8), e0202586. 10.1371/journal.pone.0202586

93. Tabachnick, K. R. (1991). Adaptation of the Hexactinellid Sponges to Deep-Sea Life. In J. Reitner & H. Keupp (Éds.), Fossil and Recent Sponges (p. 378–386). Springer. 10.1007/978-3-642-75656-6_30

94. Todd, P. A. (2008). Morphological plasticity in scleractinian corals. Biological Reviews, 83(3), 315–337. 10.1111/j.1469-185X.2008.00045.x

95. Tsakalos, J. L., Renton, M., Riviera, F., Veneklaas, E. J., Dobrowolski, M. P., & Mucina, L. (2019). Trait-based formal definition of plant functional types and functional communities in the multi-species and multi-traits context. Ecological Complexity, 40, 100787. 10.1016/j.ecocom.2019.100787

96. Untiedt, C. B., Williams, A., Althaus, F., Alderslade, P., & Clark, M. R. (2021). Identifying Black Corals and Octocorals From Deep-Sea Imagery for Ecological Assessments : Trade-Offs Between Morphology and Taxonomy. Frontiers in Marine Science, 8, 722839. 10.3389/fmars.2021.722839

97. Vlach, D. S., Pereira, O. S., Nguyen, F., Bradley, A., Mizell, K., & Levin, L. A. (2025). Megafaunal community structure on ferromanganese and phosphorite hardgrounds in the Southern California Borderland. Deep Sea Research Part II: Topical Studies in Oceanography, 224, 105560. 10.1016/j.dsr2.2025.105560

98. Watling, L., & Auster, P. J. (2021). Vulnerable Marine Ecosystems, Communities, and Indicator Species : Confusing Concepts for Conservation of Seamounts. Frontiers in Marine Science, 8, 622586. 10.3389/fmars.2021.622586

99. Wickham, H., Chang, W., & Wickham, M. H. (2016). Package ‘ggplot2’. Create elegant data visualisations using the grammar of graphics. *Version*, 2(1), 1–189. [Logiciel]. https://github.com/hadley/ggplot2

100. Williams, A., Althaus, F., Green, M., Maguire, K., Untiedt, C., Mortimer, N., Jackett, C. J., Clark, M., Bax, N., Pitcher, R., & Schlacher, T. (2020). True Size Matters for Conservation : A Robust Method to Determine the Size of Deep-Sea Coral Reefs Shows They Are Typically Small on Seamounts in the Southwest Pacific Ocean. *Frontiers in Marine Science*, A, 7, 187. 10.3389/fmars.2020.00187

101. Wong, R. HX., Monk, J., Perkins, N. R., & Barrett, N. S. (2025). Geomorphological drivers of benthic community distribution on high energy temperate mesophotic and rariphotic reefs. Marine Environmental Research, 208, 107078. 10.1016/j.marenvres.2025.107078

102. Wulff, J. L. (2006). Resistance vs recovery : Morphological strategies of coral reef sponges. Functional Ecology, 20(4), 699–708. 10.1111/j.1365-2435.2006.01143.x

103. Zawada, K. J. A., Dornelas, M., & Madin, J. S. (2019). Quantifying coral morphology. Coral Reefs, 38(6), 1281–1292. 10.1007/s00338-019-01842-4

104. Zawada, K. J. A., Madin, J. S., Baird, A. H., Bridge, T. C. L., & Dornelas, M. (2019). Morphological traits can track coral reef responses to the Anthropocene. Functional Ecology. 10.1111/1365-2435.13358

